# Merging short and stranded long reads improves transcript assembly

**DOI:** 10.1101/2022.12.13.520317

**Authors:** Amoldeep S. Kainth, Gabriela A. Haddad, Johnathon M. Hall, Alexander J. Ruthenburg

## Abstract

Long-read RNA sequencing has arisen as a counterpart to short-read sequencing, with the potential to capture full-length isoforms, albeit at the cost of lower depth. Yet this potential is not fully realized due to inherent limitations of current long-read assembly methods and underdeveloped approaches to integrate short-read data. Here, we critically compare the existing methods and develop a new integrative approach to characterize a particularly challenging pool of low-abundance long noncoding RNA (lncRNA) transcripts from short- and long-read sequencing in two distinct cell lines. Our analysis reveals severe limitations in each of the sequencing platforms. For short-read assemblies, coverage declines at transcript termini resulting in ambiguous ends, and uneven low-coverage results in segmentation of a single transcript into multiple transcripts. Conversely, long-read transcript assembly lacks strand-of-origin information and depth, culminating in erroneous assembly and quantitation of transcripts. We also discover a cDNA synthesis artifact in long-read datasets that markedly impacts the identity and quantitation of assembled transcripts. Towards remediating these problems, we develop a computational pipeline to “strand” long-read cDNA libraries that rectifies inaccurate mapping and assembly of long-read transcripts. Leveraging the strengths of each platform and our computational stranding, we present and benchmark a hybrid assembly approach that drastically increases the sensitivity and accuracy of full-length transcript assembly on the correct strand and improves detection of biological features of the transcriptome. When applied to a challenging set of under-annotated and cell-type variable lncRNA, our method resolves the segmentation problem of short-read sequencing and the depth problem of long-read sequencing, resulting in the assembly of coherent transcripts with precise 5’ and 3’ ends. Our workflow can be applied to existing datasets for superior demarcation of transcript ends and refined isoform structure, which can enable better differential gene expression analyses and molecular manipulations of transcripts.

## Introduction

Characterization of transcripts and their variants is essential to advancing our understanding of gene expression in development, responses to stimuli, and disease etiology. The advent of massively parallel short-read sequencing enabled the development of transcriptomics and discovery of biological features such as gene expression levels and pervasive genome-wide low-level transcription [1, 2]. High depth of sequencing, relatively low error rates, information about strand of origin and ability to sequence a wide range of RNA from varied upstream sources are some of the key advantages of short-read RNA-seq [3, 4]. Furthermore, multiple computational tools have been developed and benchmarked for the alignment, assembly and relative quantitation of short-read transcripts for general as well as application-specific purposes [5, 6]. However, short-read sequencing approaches bear several caveats: *i*) library preparation for short-read RNA-seq typically involves PCR amplification which is prone to length and sequence-based biases that can compromise quantification and in some cases detection [7, 8]; *ii*) the length of reads (typically 50-200 bp) is much shorter than most of the transcripts, such that transcript assembly and quantitation profoundly relies on inferential re-construction of transcripts; *iii*) differences in the computational algorithms lead to substantial inconsistencies in detection of transcripts and their splicing isoforms, variations in abundance estimation, and potential segmented assembly of a continuous transcript [6, 9-12]. These limitations are exacerbated for non-canonical transcripts such as long noncoding RNA (lncRNA) that are typically low abundance and lack canonical features of coding transcripts [9, 13].

Towards remediating these issues, new tools have become recently available. Long-read sequencing, also known as third generation sequencing, has permitted longer spans or in some cases full-length transcripts to be sequenced [14]. The MinION device from Oxford Nanopore Technologies (ONT) can directly sequence cDNA which could potentially overcome the segmentation and PCR amplification limitations of short-read sequencing [15]. While the typical read length from short-read sequencing is less than 250 nt, long reads from ONT can reach ≥10kb [16]. This can result in improved and consistent detection of transcripts and their splicing isoforms, better mapping of repeat regions, and potential mitigation of the short-read segmentation problem [17-20]. While long-read sequencing has several advantages, it too has its share of caveats, being prone to higher error rates [18, 21, 22], lower transcript coverage and depth [23, 24], lack of strand of origin information for methods such as ONT direct cDNA sequencing, and having a more limited toolkit for analysis in comparison to short-read sequencing. As many of the analysis tools were developed for DNA-level genomic assembly, they are ill-equipped to detect variations in transcript copy number and isoforms [25]. Although an array of tools has been developed for long-read RNA-seq analysis [catalogued in 26], many of them are still in their infancy and lack robustness [27]. Hybrid algorithms developed to combine short- and long-read datasets are largely built either for error correction in the long-read data [21, 28] or employ short-read as a template for long-read assembly [29]. Consequently, the output is reduced in long-read information with bias towards short-read attributes [28]. There remains a dearth of approaches that truly integrate short- and long-read datasets for transcript calling. Additionally, assembly frameworks of many of the current approaches are modeled on well-annotated high abundance canonical transcripts. Further refinement and development of long-read transcriptomic tools is warranted for analyses of non-canonical transcripts such as lncRNA which constitute a large fraction of the transcriptome [30].

LncRNA have emerged as important regulators of mammalian genomes, implicated in the regulation of genes through a diverse and growing set of mechanisms [31-36]. One class of transcripts that are highly enriched in lncRNA are the recently described chromatin-enriched RNA (cheRNA) [37-40]. CheRNA have been particularly difficult to characterize as they are highly cell-type divergent, predominantly unannotated, and so low abundance that they were largely unobserved prior to biochemical enrichment [37, 38]. Owing to these features, we reasoned that cheRNA would provide a challenging testbed to develop and benchmark long-read RNA-seq analyses for assembly and quantitation of transcripts.

In this study, we improve existing bioinformatic workflows to integrate the quantitative depth of short-read sequencing with the qualitative strengths of long-read sequencing for improved characterization of transcripts. Using short- and long-read datasets from two different cell lines with spiked-in ERCC standards, we systematically compare key parameters and biases in the read alignment and assembly of transcripts. We also report novel artifacts and further document known pitfalls in long-read datasets that can substantially impact the identity and abundance of assembled transcripts. We develop a computational pipeline, SLURP (Stranding Long Reads Using Primer sequences), to infer strand-of-origin from ONT long-read cDNA libraries (hereinafter “stranding”), markedly improving assembly of transcripts from long reads. Incorporating our stranding method, we devise a hybrid transcript assembly pipeline, TASSEL (Transcript Assembly using Short and Strand Emended Long reads), that merges qualitative features of stranded long reads with the quantitative depth of short-read sequencing. TASSEL outperforms other assembly methods in terms of sensitivity and complete assembly on the correct strand. At the molecular level, TASSEL resulted in substantially improved capture of key transcriptomic features such as transcription start and termination sites as well as better enrichment of active histone marks and RNA Pol II. To demonstrate the efficacy of TAASEL, we implement it for improved characterization of cheRNA, resolving previously ambiguous transcripts into coherent and discrete molecules. Our workflow is generally applicable to other long-read datasets with potential to refine transcript identification and isoform structure, which can enable better molecular manipulations to illuminate their mechanisms.

## Results

### Using short- and long-read RNA-seq to capture chromatin-enriched RNA

To evaluate the merits of short- and long-read sequencing platforms for assembly and molecular feature characterization of transcripts, we focused our efforts on a specific but challenging class of transcripts called cheRNA (chromatin-enriched RNA). We leveraged biochemical fractionation of nuclei [37, 41] to obtain cheRNA (Fig 1A) in two cell lines - HAP1, a near-haploid leukemia cell line derived from KBM-7 [42], and HL1, a cardiac muscle cell line derived from AT-1 mouse atrial cardiomyocytes [43]. Importantly, at the nuclear extraction step, ERCC RNA standards [44] were spiked in to allow for quality control analyses and direct comparisons between samples for both short- and long-read libraries. Fractionated RNA was then depleted of ribosomal RNA and subjected to either short-read or long-read RNA sequencing. Owing to higher quantitative depth, short-read sequencing (n=3) was done on both fractions to identify cheRNA (S1A Fig). We performed long-read sequencing (n=2) for only the chromatin fraction for comparison and conjugation with short-read cheRNA datasets. For long-read RNA-seq, RNA was polyadenylated for use with polyT primers, then subjected to library preparation using the Oxford Nanopore direct cDNA sequencing kit, and run on a MinION sequencer. We chose the cDNA sequencing platform as it has higher yield than the direct RNA sequencing platform [45]. We obtained ∼1.2M and 3M reads for HAP1 replicates, and ∼1.8M and 2.5M reads for HL1 replicates and subjected them to our analysis pipeline (S1 Table). The median sizes of reads in the long-read libraries were 1.7-1.9kb for HAP1 samples and 1.2-1.4kb for HL1 samples; replicates were nearly identical in read length profiles (Fig 1B and S1B; S1 Table). Thus, our long-read sequencing captures much longer reads than typical short-read sequencing. Next, long reads were aligned using minimap2 [46] to human (hg38) and mouse (mm10) genome assemblies for HAP1 and HL1, respectively. Aligned long reads showed strong correlation for genomic coverage between the replicates (pearson r = 0.92 and 0.86 for HAP1 and HL1, respectively) (Fig 1C and S1C). Similarly, short-read alignments also showed high correlation between replicates (S1D Fig) for both cell lines, attesting to the reproducibility of the RNA fractionation, library preparation, and alignment methodology used here.

**Figure 1.**
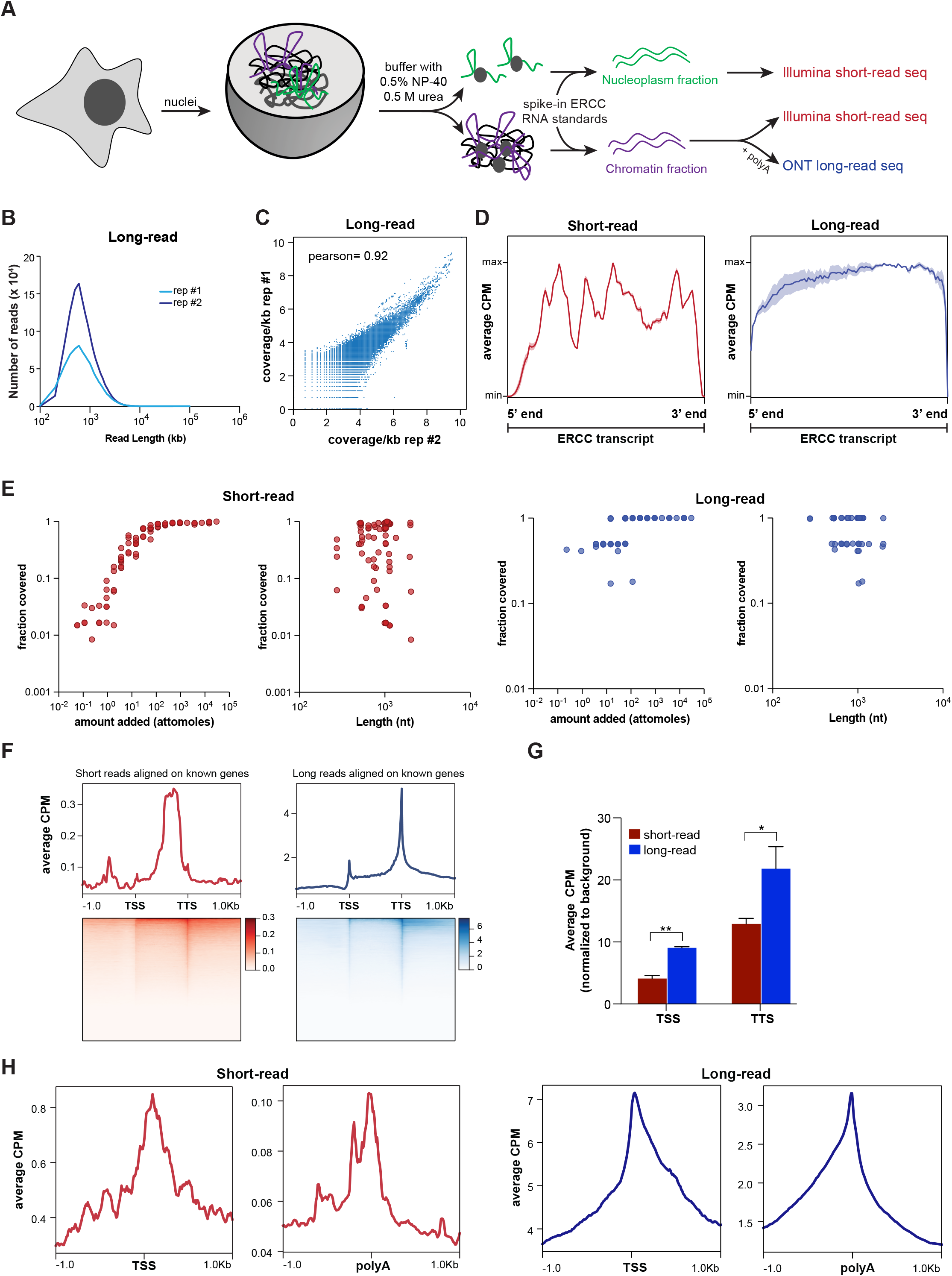
Short- and long-read RNA-seq to capture chromatin-enriched RNA. **A**. Schematic illustrating the nuclear fractionation of cells to isolate chromatin-enriched RNA (cheRNA) then subject them to short- and long-read sequencing. Nuclei from HAP1 and HL1 cells were fractionated into nucleoplasm and chromatin fractions using urea/detergent buffer [37], and ERCC standard RNA [44] were spiked in before RNA extraction and sequencing. For short-read, both the nucleoplasm and chromatin fraction RNA were subjected to single end 50 bp Illumina sequencing (n = 3). For long-read, the chromatin fraction was sequenced using Oxford Nanopore Technology (ONT; n = 2). **B**. Density plot of read lengths from long-read sequencing of HAP1 replicates (n = 2). **C**. Correlation between HAP1 replicates (n = 2) for genomic coverage of long-read alignments binned at 1 kb. Pearson correlation is shown. **D**. Average counts per million (average CPM, dark color; +/-SD, lighter color) of aligned reads across all ERCC transcripts, meta-scaled to 1000 nt, in the HAP1 short-read (n=3) and long-read (n=2) samples. **E**. Relationship between the average fraction of ERCC transcripts covered by short (left) or long (right) read alignments as a function of their amount (attomoles) or length (nt) for HAP1 samples. **F**. Metagene plots and heatmaps of mapped read coverage (average CPM) for HAP1 short- and long-read alignments, scaled to transcription start sites (TSS) and transcription termination sites (TTS) of gencode hg38v41 genes. **G**. Enrichment of TSS ± 50 bp as well as TTS ± 50 bp in short-read replicates (n = 3) and long-read replicates (n = 2) in HAP1 samples. Average counts in the indicated region for each of the replicates were normalized to a background of random 100 bp regions in the same library to account for variations in the depth of the libraries. * p < 0.05; ** p < 0.01; unpaired Student’s t-test. **H**. Metagene plots and heatmaps of mapped read coverage (average CPM) for HAP1 short (left) and long (right) reads, centered on curated transcription start sites (TSS; [49]) or termination sites (polyA; [50]).

### Spike-in standards highlight similarities and limitations in short- and long-read alignment

Transcript coverage is critical for correct assembly and estimation of abundance. The use of ERCC spike-in RNA enables us to directly compare coverage of short- and long-read alignments. Short-read alignments were highly variable in the gene body with clear lack of coverage at the 5’ and 3’ ends, whereas long-read alignments showed largely homogenous coverage across ERCC transcripts (Fig 1D and S1E). Next, we tested the extent to which a given transcript is covered as a function of its abundance or length. We observed that the fraction of a given transcript covered by short-read alignment showed a sigmoidal relationship with the amount added: near-linear sensitivity for low abundance transcripts, with coverage saturating at mid- to high-abundance transcripts (Fig 1E and S1F). While long-read alignments also showed a similar trend, they had less dynamic range, especially for low abundance transcripts. Of the 92 ERCC transcripts, 78 and 82 were detected for HAP1 and HL1 short-read datasets respectively (Fig 1E and S1F, *left*), as compared to 51 and 58 in the long-read datasets (Fig 1E and S1F, *right*). This is consistent with higher depth of the short-read datasets as compared to the long-read datasets (S1 Table). Importantly, the fraction coverage of transcripts was independent of the length of the transcripts for both short- and long-read alignments, indicating that our libraries are free of any artifactual length biases. Thus, high correlation between replicates, sensitivity to abundance, and absence of length bias demonstrate the reliability of our library-prep and alignment methods for both sequencing platforms.

### Long-read alignments are better in capturing ends of the transcripts

As short-read libraries are generated using random hexamers from fragmented RNA-samples and sequenced often with read lengths of less than 100 bp, a given read covers only a fraction of the transcript. Since long reads have the potential to capture a full-length transcript (Fig 1B and S1B), we reasoned that long-read alignment might be able to capture ends of transcripts better than short-read alignment. To formally test this hypothesis, we compared the distributions of long- and short-read alignments over known genes. While the short-read alignments showed better coverage in the gene bodies, the long-read alignments evinced better demarcation of transcription start sites and transcription termination sites of annotated genes (Fig 1F and S1G). A direct comparison showed that the 5’ and 3’ ends of a gene are more prevalent in the long-read alignments than the short-read alignments (Fig 1G). The lower gene-body coverage in the long-read alignments is consistent with previous findings [20, 47]. Additionally, we observed strong 3’-end coverage bias for long reads as reported by previous studies [20, 48].

A limitation of the foregoing analyses is that they are restricted to reference genes. However, there are many more experimentally derived TSS and TTS independent of the consensus reference annotation. To expand our analysis, we calculated the coverage of short- and long-read alignments on transcription start sites annotated on the basis of CAGE, TSS-Seq and RAMPAGE [49] as well as highly curated polyA sites [50]. Alignments from both sequencing platforms showed enrichment over these key ends of transcripts (Fig 1H and S1H). The metagene plots of long-read alignments centered over curated TSSs showed more density going into the gene body relative to upstream of the TSS for both cells lines. Conversely, long-reads centered over TTSs show more density coming from the gene body relative to downstream of the TTS for both cell lines. Corresponding short-read plots show noisier profiles and symmetrical density distributions, indicating less biologically representative capturing of 5’ and 3’ ends (Fig 1H and S1H). Thus, we conclude that long reads are capturing transcript molecule ends more effectively than short reads. Enhanced capture of intact transcripts in long-read sequencing can improve the detection of rare isoforms and gene-fusion events (see supplementary note 1). Taken together, these analyses suggest that short- and long-read data sets can complement each other for better coverage of transcriptomic elements.

### Benchmarking long-read transcript assembly

One of the crucial steps of RNA-seq analysis is assembly of transcripts from the reads as it dictates the identification of transcript structure, isoform variants, gene abundance and differential gene expression. Therefore, we compared three different transcript assembly programs: Cufflinks [51], StringTie2 [52, 56] and FLAIR [53]. For short-reads, guided assembly with StringTie2 outperformed other methods in terms of sensitivity and precision of transcript assembly (S2A Fig). Similar to the short-read data sets, guided assembly with long-read-specific StringTie2 performed better than other programs for the assembly of long reads (S2A Fig). In general, we observed that short-read transcript assembly has higher accuracy as well as precision than long-read transcript assembly. This could arise due to multiple factors such as lower depth, higher error rate and suboptimal assembly of long-read sequencing [54].

As estimation of transcript and gene abundance is one of the desired outcomes of most RNA-seq analyses, we assessed whether our long- and short-read alignment and transcript assembly are sensitive to variations in abundance and lengths of transcripts in the transcriptome. To do so, we computed abundance of ERCC spike-in standards, as their identity, abundance, and lengths are known *a priori*. We observed that the abundance of each ERCC standard in our short- and long-read libraries via StringTie assembly and quantitation showed a near-linear relationship with the concentration of each ERCC transcript added during library preparation (Fig 2A and S2B). A similar relationship was also observed using BedTools [55] which computes coverage based on alignments and is independent of transcript assembly (S2C Fig). Short-read library preparation involves PCR amplification whereas our long-read direct cDNA libraries were made without PCR. The linear range of transcript quantitation by short-read assembly and its similarity with the long-read assembly suggests that it is free of PCR biases typically thought to be associated with short-read RNA-seq analyses [7, 8]. We also observed that quantitation of short- and long-read data are independent of the length of transcripts (Fig 2A and S2B). This is particularly informative for short-read RNA-seq as it demonstrates that although individual RNA is fragmented during short-read library prep, StringTie can efficiently assemble and quantitate a wide range (0.2-2 kb) of well-annotated transcripts from short reads. Assembled transcript counts from replicates were well correlated (Fig 2B, HAP1 R^2^ =0.83; S2D Fig, HL1 R^2^= 0.74), indicating reproducibility of the extraction, sequencing, and analysis pipeline.

**Figure 2.**
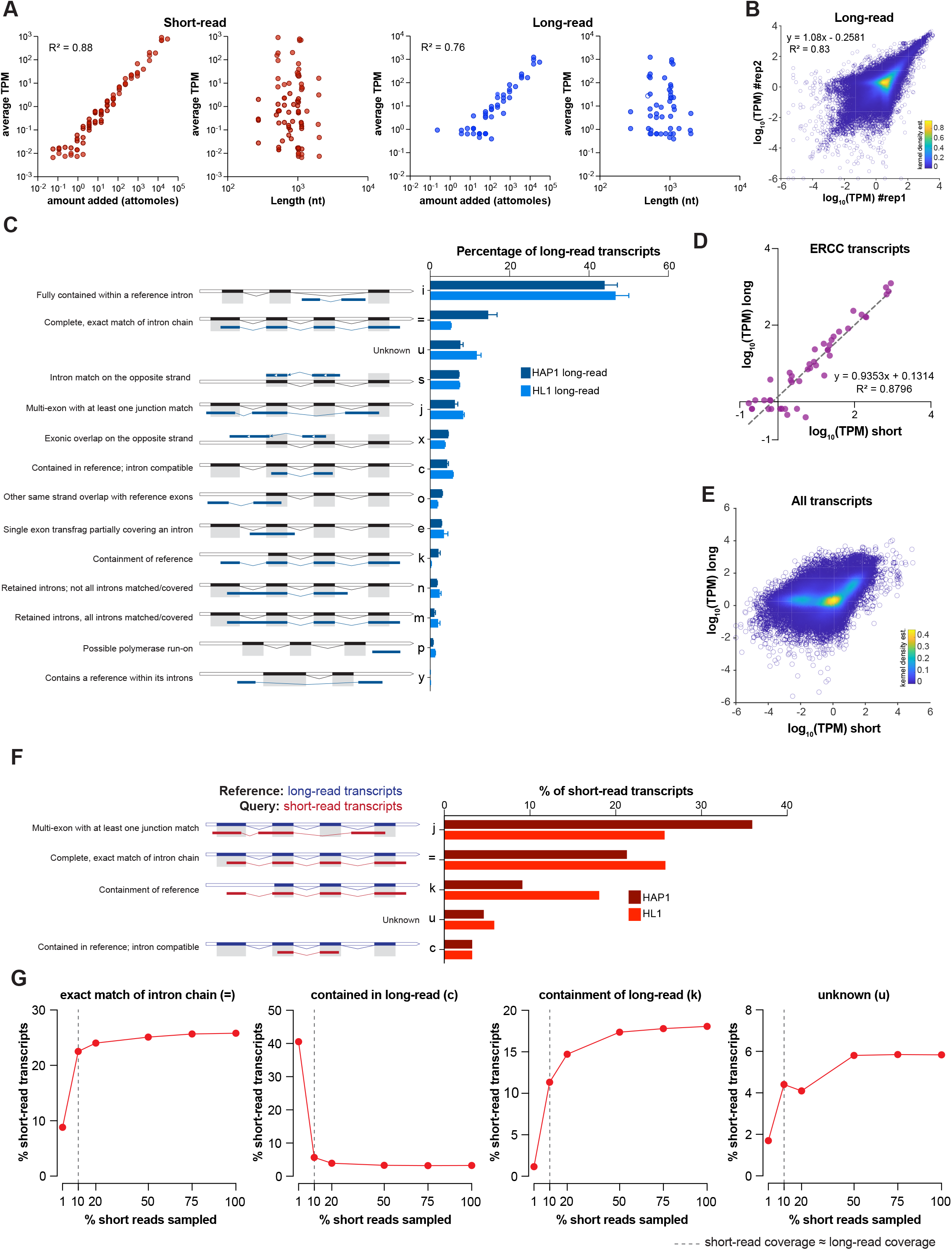
Optimization of short- and long-read transcript assembly. **A**. Average abundance (Tags Per Million; TPM) of ERCC transcripts determined by StringTie versus amount added (attomoles) or length (nt) of the transcript in HAP1 samples. (n=3 for short reads, n=2 for long reads). Short-read plots are shown in red (left), long-read plots in blue (right). R^2^, correlation coefficient for linear regression. **B**. Correlation between HAP1 replicates for long-read transcriptome assembly by StringTie. Shown are the scatter plots of abundance (TPM) of each transcript in the two replicates. Colors correspond to kernel density estimations of scatter plot distribution. R^2^, correlation coefficient for linear regression. **C**. Comparison of structure of transcripts assembled by StringTie for HAP1 and HL1 long-read samples with structure of reference genome transcripts. The class codes for relationship between the assembled transcript and the closest reference transcript were deduced from gffcompare. **D**. Correlation plots for the average abundance (TPM) of ERCC spike-in transcripts in the short-read datasets (n=3) with the average abundance in the long-read datasets (n=2) of HAP1 samples, determined by StringTie. Shown are the transcripts detected by both sequencing platforms. **E**. Scatter plots comparing the average abundance (TPM) of all transcripts in the short-read datasets (n=3) and the long-read datasets (n=2) in HAP1 samples. Abundance was calculated using the re-estimation function of StringTie from the merged transcriptome. Color bars correspond to kernel density estimations of scatter plot distribution. **F**. Structural comparison of short-read assembled transcripts with the long-read assembled transcripts for HAP1 and HL1 samples. The long-read transcriptome assembly was used as the reference and short-read as the query in gffcompare. **G**. Percent of transcripts in the down-sampled short-read dataset for several key transcript structures (class codes in panel 2F) using long-read transcripts as a reference in HL1 samples. Note that depth of 10% of HL1 short-read dataset (dotted line) is approximately equivalent to the depth of long-read dataset.

Having established the efficacy of StringTie in the assembly of transcripts from long reads, we compared the assembled long-read transcriptomes with reference transcriptomes (Fig 2C). We observed that more than 80% of long-read transcripts overlapped with the reference transcriptome: 5-14% of transcripts matched exactly with the reference transcriptome, while 6-8% showed a match of at least one intron-exon junction. In contrast, 45-60% of transcripts assembled from short-read samples showed an exact match with the reference transcriptome (S2E Fig). Similar to previous analyses [56], we observed that a large percentage of the long-read assembled transcripts (>40%) were fully contained in reference gene introns. The prevalence of such transcripts is higher in our dataset, possibly because of the enrichment of nascent transcripts in the chromatin-bound transcripts [57-59]. Notably, many transcripts in our long-read datasets (∼7%; Fig 2C class code “s”), in contrast to the short-read datasets (S2E Fig, class code “s”), were assembled to the opposite strand of the reference transcriptome. This is because strand information is lost during the preparation of cDNA for ONT direct cDNA long-read RNA-seq. In contrast, the short-read libraries were prepared using a method that preserves the strand information.

### Comparison of short- and long-read data

After establishing that our long-read datasets pass initial quality control measures, we assessed long-read sequencing performance relative to the short-read sequencing. To do so, we leveraged matching short-read datasets for each cell line, which also contained spiked in ERCC RNA. Analysis of averaged TPMs of ERCC standards from StringTie for long reads (n=2) against short reads (n=3) showed strong correlation between the HAP1 and HL1 datasets (R^2^= 0.88 and 0.91, respectively) (Fig 2D and S2F). Expanding this analysis to all transcripts assembled by StringTie showed high correlation of quantitation for abundant transcripts (Fig 2E and S2G). However, we observed less correlation for low abundance transcripts (log_10_(TPM) < 0), as short-read sequencing showed more dynamic range than long-read sequencing for these transcripts. While both transcript assembly platforms peaked at similar lengths in their distributions, short-read assemblies had a higher number of long transcripts (S2H Fig).

Then, we compared the structure of short- and long-read assembled transcripts. More than 20% of long-read transcripts in the HAP1 dataset and more than 25% of long-read transcripts in the HL1 dataset showed an exact match with their short-read counterparts (Fig 2F). Additionally, ∼30% of multi-exon long-read transcripts had at least one intron-exon junction matched with the corresponding short-read transcripts. Taken together, these two features suggest high convergence of the two platforms. The differences in transcript structure between the two platforms could arise due to differences in the depth of sequencing or the inherent differences in the sequencing and assembly methodologies. To test the effect of depth differences, we compared the long-read transcriptome structure with that derived from down-sampled short reads (Fig 2G). The percentage of short-read transcripts matching exactly with a long-read transcript increases concomitantly with the depth of short-read sequencing (Fig 2G, class code “=”). The inverse trend is seen for short-read transcripts that are contained within the long-read transcripts (Fig 2G, class code “=”), emphasizing the fact that assembly from low-depth short-read RNA-seq leads to artifactual segmentation of transcripts. Conversely, we detected that the incidence of long-read transcripts contained within short-read transcripts increases with increase in the depth of short-read sequencing (Fig 2G, class code “k”). Likewise, the proportion of short-read transcripts with no counterpart in the long-read dataset was higher in the high-depth short-read dataset (Fig 2G, class code “u”). These analyses indicate that sequencing depth plays a predominant role in limiting concordant assembly of transcripts between the short- and long-read platforms.

Our analyses highlight two main caveats of transcript assembly by long reads: assembly of long-read transcripts on the incorrect strand due to the lack of strand of origin information, and containment of long-read transcripts in the reference genes due to the lack of sequencing depth. We undertook to develop a computational pipeline to overcome these limitations.

### Development of computational stranding of long read reads to improve transcript assembly

There are many methods to prepare and sequence stranded libraries for short-read RNA-seq [60] which culminate in accurate assembly and analyses of transcripts [61]. However, no standard method exists for either the preparation or the computation of stranded libraries for ONT direct cDNA long-read RNA-seq. Consequently, long-read RNA-seq suffers from inaccurate alignment of reads to the incorrect strand of the locus of origin, leading to assembly of transcripts opposite to their true strand (Fig 2C). As the existing software for stranding of long reads yielded only 6% of stranded reads (S3A Fig), we developed a computational “stranding” pipeline, SLURP (Stranding Long Reads Using Primer sequences (S3D Fig)). Our method exploits the sequences of primers used during preparation of cDNA long-read libraries to infer strand of origin (Fig 3A). Specifically, primer 1 (with a polyT 3’ end) is used to synthesize the first strand of cDNA by reverse transcriptase, which subsequently switches its template strand using primer 2 (with a GGG 3’ end) to synthesize the second cDNA strand. Therefore, we reasoned that the presence of primer 1 or the reverse complement of primer 2 in a read would indicate that it originates from the first strand, whereas the presence of primer 2 or the reverse complement of primer 1 would denote that the read is derived from the second strand. As the primers are incorporated in the majority of the reads, we took advantage of a motif-enrichment tool, MEME [62], to predict the sequences of primer 1 and primer 2 (S3B Fig). We checked the location and prevalence of the two primer sequences in the long reads. Consistent with priming from the transcript termini, primer 1 and primer 2 were highly enriched within first 100 bp of the reads while their reverse complements were enriched in the last 100 bp of the reads (Fig 3B). To our surprise, we observed that the prevalence of primer 1 (>53% of reads) is considerably higher than primer 2 (<9%), suggesting that the first strand synthesis step is more efficient than strand-switching and second strand synthesis. This skew may account for the coverage bias at the 3’-end of long-read RNA-seq reported in many previous studies [reviewed in 20].

**Figure 3.**
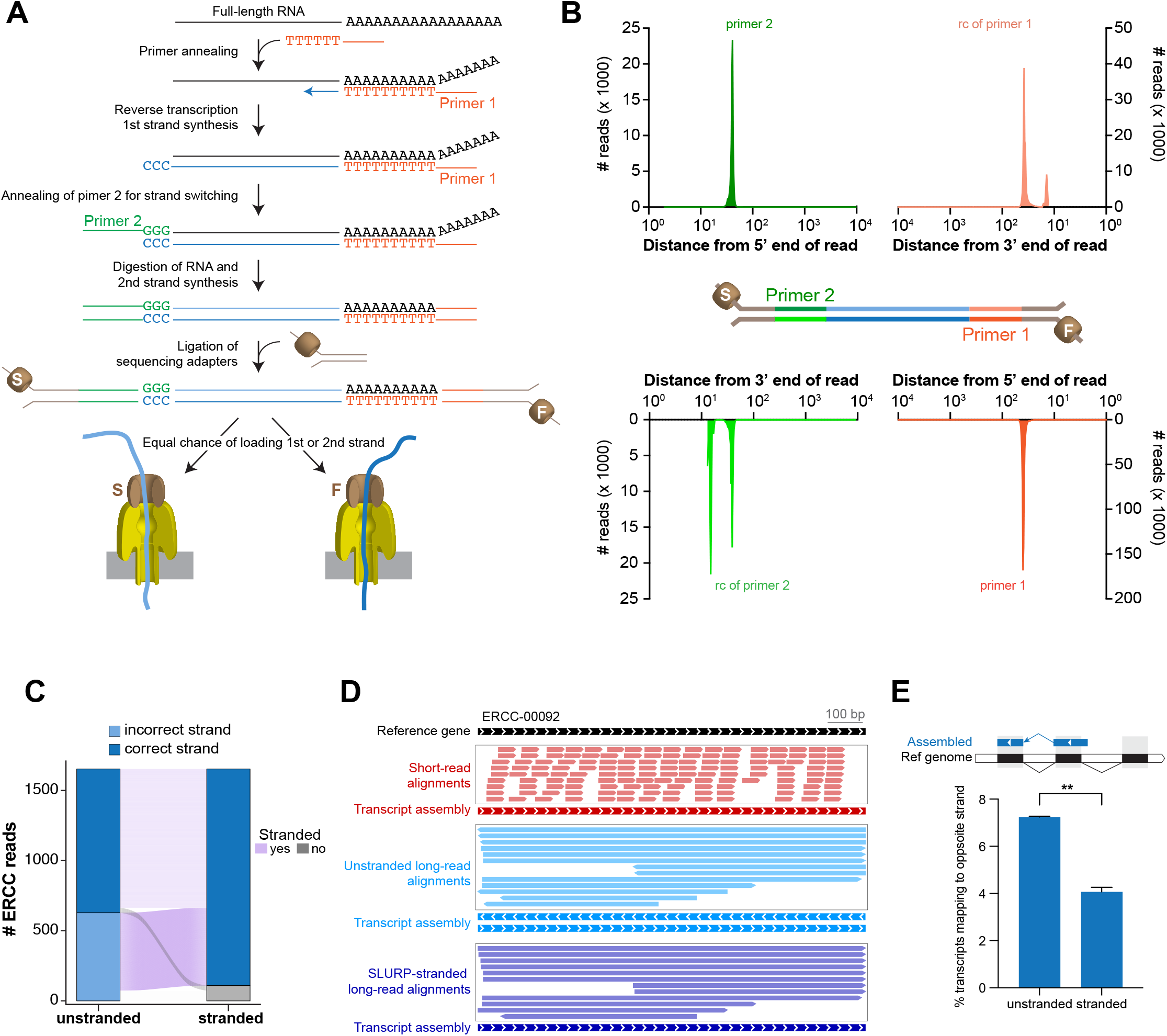
Primer-based elucidation of the strand of origin of long reads. **A**. Schematic of ONT long-read cDNA library preparation. Primer 1 with 3’ polyT initiates first strand synthesis by reverse transcriptase that adds non-templated -CCC-at the end. Primer 2 with -GGG-end anneals to the first strand and mediates strand switching for synthesis of the second strand. Sequencing adapters are subsequently ligated to the double stranded cDNA molecule. Either of the two strands (F: first, S: second) can be loaded in a given sequencing nanopore. **B**. Location and enrichment of sequencing primers (primer 1 and primer 2) and their reverse complements in the long reads. **C**. Change in the correct strand mapping of unstranded and stranded long-reads to ERCC transcripts. **D**. Assembly of ERCC-00092 transcript by unstranded and stranded long reads. **E**. Comparison of percentage of transcripts mapping to the opposite strand of reference transcripts in the unstranded and stranded long-read transcriptome of HAP1 cells (n=2; mean ±SD; ** p < 0.01 unpaired Student’s t-test).

We compared different combinations of primer searches as well as permissible number of mismatches to strand our long-read libraries. Guided by their prevalence (Fig 3B), we restricted our search of primer sequences to the first or last 100 bp of reads to minimize artifactual stranding due to coincidental occurrence of these sequences in read interiors. We observed that a 3-criteria (primer 1, primer 2 and reverse complement of primer 2) search combined with 2 mismatches provided an appropriate balance between the number of assembled transcripts and extent of correct stranding with respect to the reference transcriptome (S3C Fig). We checked the efficacy of SLURP on ERCC spike-in standards, as their precise strand of origin is known *a priori*. SLURP successfully reassigned the majority (>90%) of the long reads originated from ERCC loci to their correct strand (Fig 3C). Importantly, while the alignment of unstranded long reads led to ambiguous assembly of two ERCC-00092 transcripts in opposite directions, alignment of stranded long reads led to assembly of one correct transcript (Fig 3D). We expanded our analyses to all assembled transcripts and observed that the stranding of long reads also led to significant reduction of transcripts assembled to the opposite strand of the reference transcriptome (Fig 3E). Transcripts that are still assembled on the opposite strand could be a result of a residual pool of reads that were not stranded or *bona fide* anti-sense transcripts [63]. As a proof of general application, we tested SLURP on publicly available long-read data [47]. We were able to substantially reduce the mapping of long reads to the incorrect strand (S3E Fig), demonstrating that our stranding pipeline can be used as a general tool to infer strand information from long-read cDNA data which increases the accuracy of transcript assembly and any subsequent analysis.

### Long-read libraries are prone to reverse transcriptase cDNA synthesis artifacts

As per the library preparation method, primer 1 initiates the synthesis of the first strand and primer 2 initiates the synthesis of the second strand after the reverse transcriptase switches strands (Fig 3A). Thus, the occurrence of primer 1 and its reverse complement should be mutually exclusive. Surprisingly, we discovered that many long reads have the primer 1 sequence in their first 100 bp and the reverse complement of primer 1 in their last 100 bp (Fig 4A). The occurrence of the primer 1 sequence at each end of a single read would not be anticipated from the way the library was prepared, yet this is observed in ∼6% of total reads (Fig 4B). To test the prevalence of this artifact, we probed a published long-read data set [47] and detected similar levels of such reads (Fig 4B). Further examination revealed that these reads map to the same genomic locus twice in opposite directions (Fig 4C). Notably, we found that one of these two alignments is flagged as a “supplementary alignment” by the minimap2 aligner while the other is considered as the primary alignment. We reasoned that such an alignment could be possible if the reads are palindromic. Indeed, secondary-structure prediction shows that most of these reads are largely palindromic within base calling error, with one half of the read being the reverse complement of the other (Fig 4D, 4E and S4A). Surprisingly, filtering of supplementary alignments led to substantial changes in the computation of coverage and abundance of transcripts (Fig 4F). We speculate that these reads originate when the reverse transcriptase switches strands due to micro-homology [64] and continues to synthesize cDNA using the first strand as the template (Fig 4G), leading to artifactual doubling of read length via reverse complement synthesis. Consistent with this notion, we detected virtually no palindromic reads in the direct RNA long-read libraries that do not use reverse transcriptase for cDNA synthesis (Fig 4B).

**Figure 4.**
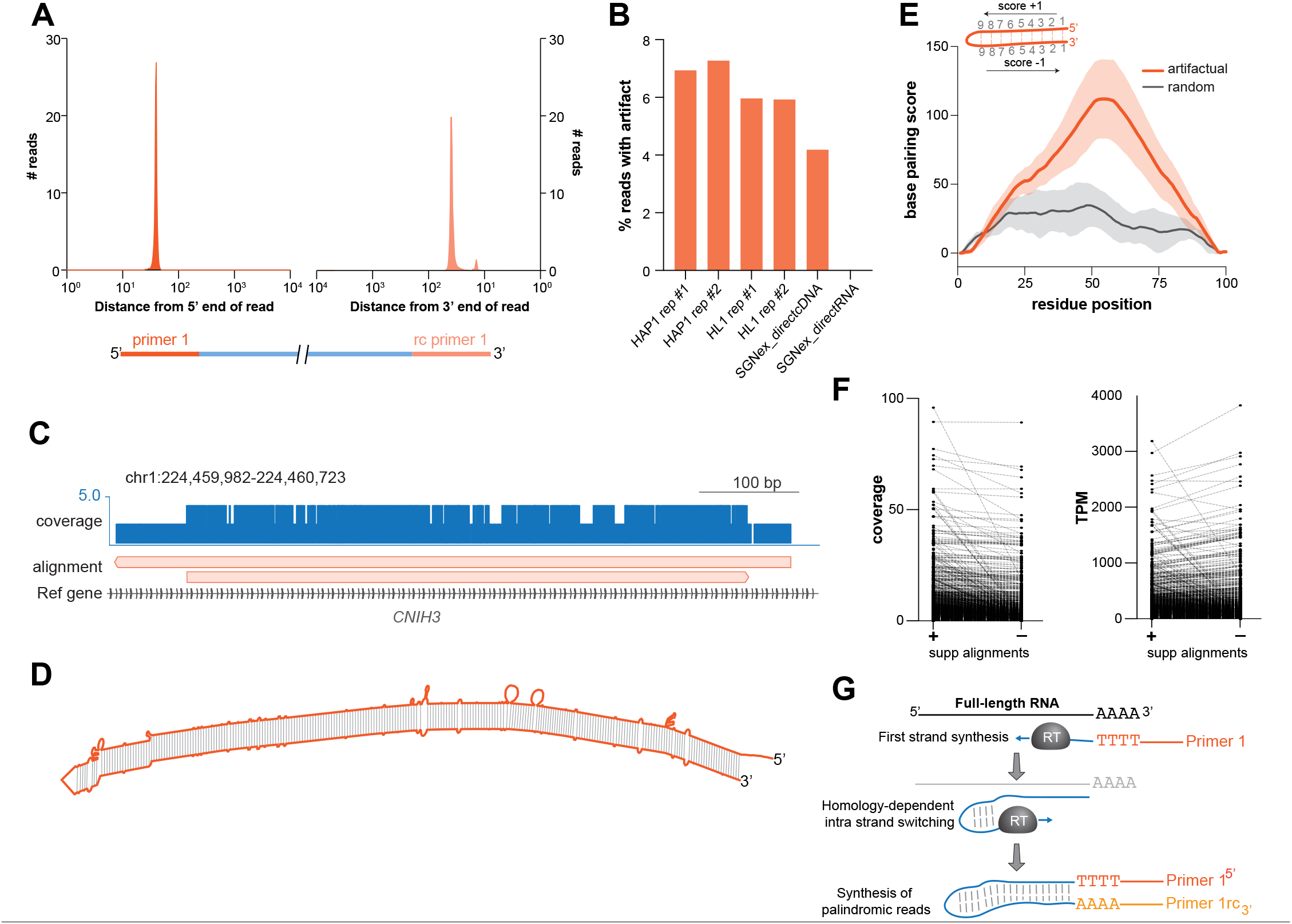
Widespread cDNA synthesis artifact in long reads. **A**. Plot showing enrichment of primer 1 at the 5’ end and the reverse complement of primer 1 at the 3’ end of artifactual reads. **B**. Prevalence of artifactual reads with primer 1 as well as its reverse complement in the libraries generated in the current as well as previously published studies. Note the lack of such artifactual reads in the library generated from direct RNA sequencing. **C**. Genome browser track showing the alignment of an artifactual read twice at the same locus (*CNIH3*) in the opposite directions. Note that this is the only read that mapped to this locus; depth of coverage by minimap2 alignment is doubled in the places where the read maps twice. **D**. Secondary structure prediction of the read in panel **C** by RNA-fold shows that it is largely palindromic. **E**. Meta mountain plot of the extent of palindromes in the artifactual (orange, n=50) and the same number of randomly selected (black) reads from the whole set (not restricted to non-semi-palindromic). Base pairing score of the given base from RNA fold structure is increased by 1 if the base is paired downstream and decreased by 1 if the base is paired upstream; no change for an unpaired base. Scores were scaled to 100 bp meta-transcript. Shaded region indicates SD. **F**. Measurement of the coverage (left) and abundance (TPM; right) of transcripts by StringTie before and after filtering the supplementary alignments. **G**. Model depicting a possible source of artifactual reads in the long-read libraries. Primer 1 initiates synthesis of the first strand by the reverse transcriptase (RT). A micro-homology region in a read may create a short hairpin which would lead to switching of the template strand from the original RNA molecule to the first strand which is being synthesized, producing a continuous palindromic read.

Hence, these analyses reveal a widespread heretofore unknown artifact in ONT long-read cDNA libraries that contribute to error in read length and counting. The mapping error by minimap2 is propagated to transcript assembly and quantitation by StringTie, which could negatively impact the accuracy of gene expression analyses.

### Integration of short-read data with stranded long-read data improves transcript assembly

To overcome the limitations of long-read transcript assembly, we developed a hybrid transcript assembly pipeline, TASSEL (Transcript Assembly using Short and Strand Emended Long reads), that incorporates long range information of stranded long reads with high depth of short-read sequencing, with very low additional computational burden (Fig 5A). First, we compared TASSEL to other hybrid and standalone long-read transcript assembly programs for transcript assembly of ERCC spike-in standards (Fig 5B). FLAIR [53], Bambu [65] and StringTie Mix [29] failed to assemble ∼40 of the 92 ERCC transcripts, while only 13 transcripts were not assembled via TASSEL. FLAIR and StringTie Mix also assembled many incomplete ERCC transcripts where the assembled transcripts were shorter than the actual molecular standards. Although FLAIR involves a “correction” step, it did not correct for mapping and assembly of long read transcripts on the incorrect strand. No such assembly on the incorrect strand was detected for StringTie Mix, Bambu and TASSEL. Most importantly, TASSEL, by far, assembled the highest number of complete ERCC transcripts on the correct strand. When tested as a function of transcript abundance, we observed a large variation in the rate of false negative assembly of ERCC transcripts amongst the tested assembly methods (Fig 5C). TASSEL showed a substantially lower false negative rate than other methods, indicating its higher sensitivity. A key parameter of assembling the correct and coherent transcript is determination of its ends. The ends of TASSEL-derived ERCC transcripts matched completely with the actual ends of these standard transcripts (Fig 5D and S5A). On the other hand, many transcripts assembled with StringTie Mix were artifactually truncated at their 5’ and/or 3’ ends.

**Figure 5.**
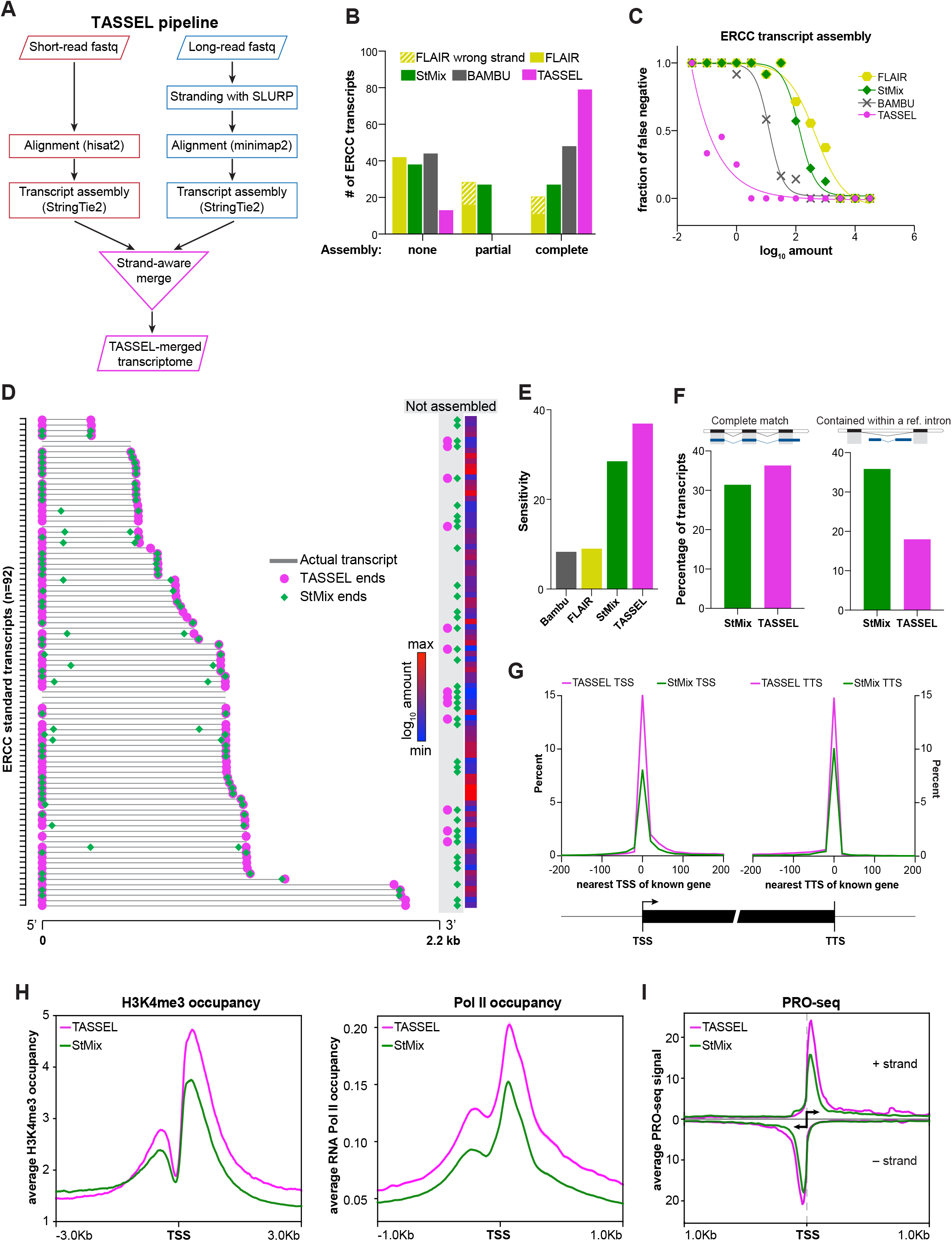
Improved transcript assembly with strand-aware hybrid pipeline. Workflow of TASSEL (Transcript Assembly using Short and Strand-Emended Long reads) pipeline. Transcripts obtained from short-read RNA-seq are merged with those obtained from stranded long reads in a strand-aware manner. Number and extent of ERCC standard transcripts assembled by the indicated assembly method in the HAP1 dataset. StMix, StringTie Mix. **C**. False negative rate of assembling ERCC transcripts, by each of the indicated assembly methods, as a function of their abundance in the HAP1 dataset. **D**. Ends of the 92 ERCC transcripts (arranged in the increasing order of length) assembled by TASSEL (magenta circle) and StringTie Mix (StMix, green diamond) in the HAP1 dataset. Gray bar indicates actual transcript. The color bar indicates the abundance of the given transcript. **E**. Sensitivity of the indicated assembly methods at the locus level. Transcriptome assembled by the given method in the HAP1 dataset was compared against reference annotation (gencode hg38v35) using gffcompare. **F**.Percent of assembled transcripts that match completely with a transcript (left) or contained within an intron (right) of reference transcript (gencode hg38v35), using StringTie Mix or TASSEL in the HAP1 dataset. **G**. Proximity of TSS (left) and TTS (right) of known genes (gencode hg38v41)to the TSS and TTS of transcripts assembled by TASSEL or StringTie Mix in the HAP1 dataset. **H**. Enrichment of H3K4me3 (left, normalized to input) and RNA Pol II (right, normalized to the total number of mapped reads) at the TSS of transcripts assembled by TASSEL or StringTie Mix in the HAP1 dataset. H3K4me3 and RNA Pol II occupancy calculated from ChIP-seq data from HAP1 cells [90, 96]. **I**. Average PRO-seq signal at TSS of transcripts assembled by TASSEL or StringTie Mix on the positive (top) and negative (bottom) strands. Normalized PRO-seq data in HAP1 cells were obtained from [92].

Next, we compared the entire transcriptome assembled by these programs with the reference transcripts. Although Bambu performed better than StringTie Mix for ERCC assembly, it had the lowest sensitivity of assembly at the locus level (Fig 5E). Here again, TASSEL outperformed the other tested assembly programs. As StringTie Mix was closest to TASSEL in performance, we performed further comparisons between the two. In comparison to StringTie Mix, TASSEL showed a modest increase in transcripts that match completely to reference transcripts (Fig 5F and S5B, left). Importantly, there was a substantial decrease in the assembled transcripts that are contained within the reference transcripts (Fig 5F and S5B, right). We reason that this improvement in TASSEL is due to the incorporation of higher depth of short-read sequencing.

We also tested how TASSEL compares to StringTie Mix in the accurate determination of transcription start sites (TSS) and transcription termination sites (TTS) of all assembled transcripts. For this, we compared the TSS and TTS of TASSEL and StringTie Mix transcripts with those of known genes (Fig 5G). TASSEL showed much better overlap with known TSS and TTS than StringTie Mix, indicating that TASSEL assembles more coherent transcripts than StringTie Mix. To further evaluate the accuracy of the assembled 5’-ends, agnostic of the reference assemblies that may not reflect TSS in a particular lineage [66], we sought to compare TASSEL and StringTie Mix for enrichment of other genomic features of 5’ ends from the same cell line. At the molecular level, transcription start sites are sites of high H3K4me3 deposition (epigenetic mark of active TSS [67]) and RNA Polymerase II (RNA Pol II) occupancy [68]. TSS derived from TASSEL were more enriched for these two functional entities than those derived from StringTie Mix (Fig 5H, S5C). Precision Run-On sequencing (PRO-seq), used to map the location of active RNA polymerase, can provide estimates of the 5’ end of transcripts [69]. TASSEL-derived TSS showed substantially higher PRO-seq signal than StringTie Mix-derived TSS (Fig 5I), suggesting that TASSEL TSS are better indicator of active TSS than StringTie Mix TSS. Collectively, our data show that TASSEL outperforms contemporary transcript assembly methods for correct and complete assembly of transcripts as well as enrichment of transcriptionally relevant features. Thus, we felt confident moving forward to use our method to interrogate assembly of cheRNA as a challenging testbed.

### TASSEL improves cheRNA identification

Based on the foregoing analyses, we reasoned that TASSEL can be used to improve the assembly of cheRNA which have been challenging to characterize due to their non-canonical features, low abundance, and high variance. First, we used short-read RNA-seq on the chromatin and nucleoplasm fractions to identify cheRNA as it provides higher depth and dynamic range. The control ERCC spike-in standards were detected at equivalent levels in the two fractions (S6A Fig). We defined cheRNA based on their marked enrichment (>4 fold) in the chromatin fraction relative to the nucleoplasm fraction (Fig 6A). Using differential expression analyses, we detected 6,746 and 4,969 cheRNA genes in HAP1 and HL1 cells, respectively. The detected cheRNA genes were proximal to cell line-specific protein coding genes (S6B Fig), suggesting potential biological significance.

The exact transcript structure of the detected cheRNA is not clear due to caveats in short-read RNA-seq, such as lower coverage of transcript start and end sites (Fig 1F, 1G and S1G) and apparent segmentation of transcripts due to the short length of aligned reads. When compared to long-read dataset, we observed moderate to high enrichment of long reads across cheRNA genes in comparison to their upstream and downstream regions (Fig 6B), suggesting that cheRNA genes, identified through short-read RNA-seq, are represented in the long-read datasets. Then we compared the gene structure of cheRNA genes from the short-read datasets with the transcriptome assembled from the long-read datasets. More than 40% of cheRNA genes from the HAP1 short-read dataset and more than 20% from the HL1 short-read dataset had a counterpart in the long-read datasets with more than 90% overlap (Fig 6C). We observed that cheRNA genes with minimum overlap were significantly less abundant than those with maximum overlap (S6C Fig). While this analysis shows that a large fraction of cheRNA gene structures inferred from short-read analysis match that of the long-read assembly, it also reveals that long reads fail to assemble coherent cheRNA transcripts in regions of low coverage. An example is highlighted in Fig 6D, where a cheRNA locus is detected on the basis of high enrichment in the chromatin fraction as compared to the nucleoplasm fraction in the short-read data. Although the coverage track indicates a single gene, StringTie assembly of short reads predicts two cheRNA genes at this locus. The StringTie assembly of the original long reads resulted in assembly of three transcripts – one on the Crick strand and two on the Watson strand. StringTie mix [29] also fails to predict the correct transcript. Strikingly, use of TASSEL led to the assembly of one transcript in the correct direction. TASSEL not only assembled the transcript on the correct strand but also resolved the segmentation problem. We computed that ∼30% of the original cheRNA were segmented prior to application of TASSEL (Fig 6E). To test that this computational approach is biologically meaningful, we compared RNA Pol II occupancy at the TSS of originally segmented or unsegmented cheRNA genes with TASSEL-refined cheRNA genes. Consistent with the expectation of Pol II enrichment decorating actively transcribed promoter regions [68], we detected substantial increase in Pol II enrichment at the TSS of refined cheRNA (Fig 6F and S6D). There were also better indications of divergent transcription – a characteristic of *bona fide* promoters – from the TSS of refined cheRNA genes. Additionally, there was marked improvement in coverage of long reads over refined cheRNA genes with sharpened demarcation of 5’ and 3’ ends (Fig 6G vs Fig 6B). These analyses highlight the utility of TASSEL for improved assembly of correct transcripts and genes, even for one of the most challenging classes of molecules.

**Figure 6.**
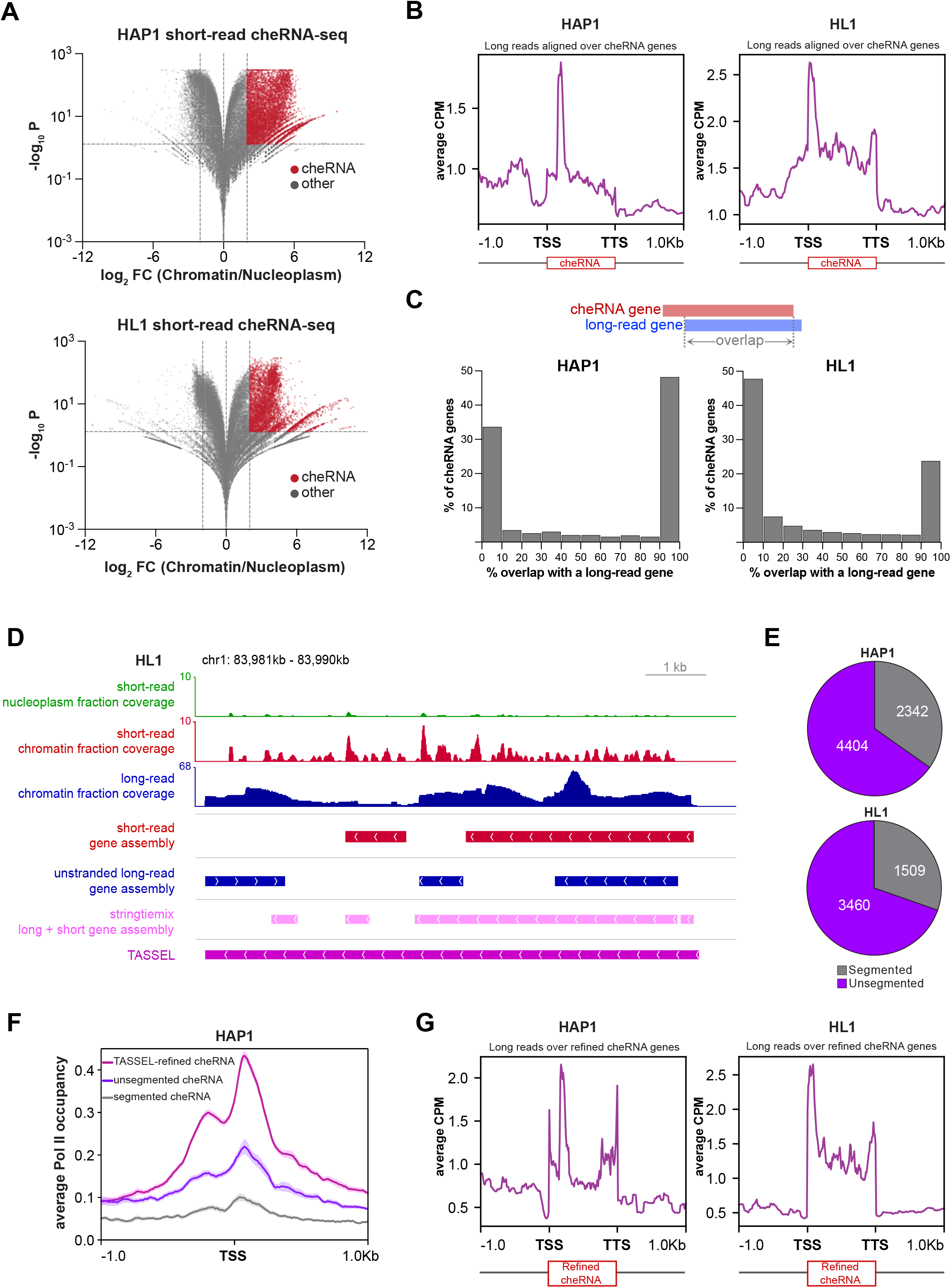
Improved cheRNA characterization by integration of short-read assemblies with stranded long-read assemblies. **A**. Volcano plot for enrichment of genes in the chromatin fraction vs adjusted *p* value. DESeq2 on short-read counts by StringTie was used to obtain cheRNA genes (red) that are significantly (Benjamini and Hochberg adjusted *p* < 0.05) enriched (>4 fold) in the chromatin fraction as compared to the nucleoplasm fraction. **B**. Metagene plots and heatmaps of mapped read coverage (average CPM) for HAP1 (left) and HL1 (right) long-read alignments, scaled to the cheRNA genes in the corresponding samples. **C**. Histogram depicting the extent of overlap between cheRNA genes from short-read assembly and genes assembled from long-read assembly in HAP1 and HL1 samples. **D**. Example cheRNA locus depicting the efficacy of the optimized methodology and improved assembly of cheRNA genes. Top: coverage from short-read nucleoplasm and chromatin fractions shows high enrichment in the chromatin fraction. Bottom: assembly by indicated methods shows resolution of the segmentation problem by merging short-read with the stranded long-read assembly. **E**. The number of segmented cheRNA genes identified in HAP1 and HL1 short-read datasets after merging of short-read and stranded long-read transcriptome. **F**. Metagene plot depicting average RNA Pol II occupancy (solid lines; shaded region indicates SE) at the TSS (± 1 kb) of segmented, unsegmented and TASSEL-refined cheRNA genes in HAP1 dataset. RNA Pol II occupancy calculated from ChIP-seq data from HAP1 cells [90]. **G**. Metagene plots and heatmaps of mapped read coverage (average CPM) for HAP1 (left) and HL1 (right) long-read alignments, scaled to the TASSEL-refined cheRNA genes.

## Discussion

With the increased interest in long-read RNA-seq, current efforts are directed towards weighing the merits of long-read RNA-seq over its short-read counterpart as well as development of new tools for improved coalescence of the two platforms [20, 29, 47, 70, 71]. To examine whether short- or long-read RNA-seq data or a combination of both would permit robust transcript characterization, we compared key features of read alignment, read coverage, transcript assembly, and capture of biologically relevant attributes of transcripts from short- and long-read RNA-seq. To sharpen the challenge, we focused on chromatin-enriched RNA, a class of largely unannotated, low-abundance transcripts that have presented problems for previous short read analyses [37, 38]. We performed Illumina short-read and ONT direct cDNA long-read RNA-seq in parallel on multiple biological replicates in cell lines of mouse and human origin. Consistent with the performance of the two platforms, we obtained 2-6X coverage/bp for short-read sequencing and 0.5-0.7X for long-read sequencing. Alignments of reads from both platforms to the corresponding genomes were highly correlated between replicates, attesting to their reproducibility. Our use of spike-in ERCC RNA standards [44] enabled us to objectively compare the performance of the two platforms. Both showed a near-linear relationship for observed versus expected transcript abundance, validating their use in transcript quantitation and differential gene expression analyses. Owing to its higher sequencing depth, the short-read platform had a greater dynamic range for transcript detection, fraction of transcript covered as well as quantitation. In contrast, long-read coverage was more homogenous across ERCC transcripts and provided better coverage of 5’ and 3’ ends than short-read alignments. Collectively, our data suggest that short-read RNA-seq has quantitative advantages whereas the long-read RNA-seq has enhanced qualitative attributes. A hybrid approach that retains the best aspects of each can result in more definitive transcriptome characterization.

### Strengths and weaknesses of short- and long-read RNA-seq

As transcript assembly is critical for most of the downstream analyses, we systematically compared multiple transcript assembly approaches. Consistent with the previous studies [52, 56], reference-guided assembly with StringTie2 showed higher sensitivity as well as precision for transcript assembly for both short- and long-read RNA-seq, relative to other tested programs. A large fraction of long-read transcripts was fully contained within the reference intron, perhaps largely due to the enrichment of nascent transcripts in chromatin fractions of nuclear RNA [57-59]. We observed a high overlap between transcripts assembled in the short- and long-read datasets. A direct comparison of the structure of transcripts showed that many in the long-read datasets were “contained” in the short-read transcripts, largely due to the higher depth of the short-read sequencing. Therefore, lower depth of long-read sequencing can cause truncation of assembled transcripts at low coverage regions. Consistent with this notion, we observed a higher abundance of longer transcripts in the short-read assembly. The lower depth of long-read assembly also impacted the quantitation of low abundance transcripts when compared to the short-read transcriptome, cautioning its use for pan-transcriptome quantitation on its own.

### SLURP strands and improves transcript assembly

The lack of strand of origin information in the ONT direct cDNA long-read sequencing hinders direct comparison to reference transcriptomes and short-read assemblies. Consistent with this limitation, we report clear evidence of mapping of ∼40% of long reads to the incorrect strand and consequent ambiguous assembly of transcripts. To address this problem, we developed and benchmarked an accessible computational stranding pipeline, SLURP. Using SLURP, we corrected the erroneous mapping of long reads, resulting in a striking improvement in the assembly of spiked-in ERCC standard transcripts and assembly of transcriptomes. StringTie Mix [29], a hybrid assembly program, permitted comparable stranding efficiency but resulted in truncated transcripts, seemingly due to overweighting of short reads in the hybrid assembly. Unlike StringTie Mix, SLURP can be used as a general-purpose standalone pipeline without the corresponding short-read data. Importantly, our stranding pipeline can be broadly applied to preexisting long-read RNA-seq studies for improved transcript assembly, comparison and integration with short-read RNA-seq data, and high-confidence detection of *bona fide* antisense transcripts shown to play a critical role in gene regulation [72].

### Prevalent artifacts in direct ONT cDNA long-read sequencing

Our in-depth stranding analysis led to the serendipitous discovery that long-read ONT cDNA libraries are plagued with a cDNA synthesis artifact. Surprisingly, ∼6% of long-read cDNA libraries are palindromes (within base-calling error). As we did not detect such reads in the ONT direct RNA library, we attribute these reads to an artifact of first strand synthesis, wherein reverse transcriptase switches from its initial RNA template to its product cDNA strand for further elongation. This finding is distinct from previously reported reverse transcriptase artifacts such as artificial splicing, in which the reverse transcriptase jumps from one direct repeat to another direct repeat of the same RNA or different RNA molecule, causing the deletion of the intervening sequence [73, 74]. There is less chance of formation and detection of such palindromes in the short-read datasets as short-read libraries are synthesized from fragmented RNA and sequenced with shorter read lengths, often with enzymes that lack template switching activity. Alignment using standard long-read aligners such as minimap2 leads to double mapping of these reads to the same genomic region and flagging of one of the two alignments as a supplementary alignment. However, such an alignment not only inflates coverage statistics at the alignment stage but also leads to imprecise assembly and quantitation of all transcripts. It is important for the field to be aware of this likely prevalent artifact in long-read cDNA libraries which, to our knowledge, has not yet been reported. In principle, alternate cDNA library preparations with non-template switching reverse transcriptases or future aligners and assembly algorithms adapted to resolve these palindromes can be employed to accommodate these pitfalls.

### Enhanced detection of molecular features promises to improve functional analyses

Knowledge of transcript ends is critical for downstream experimental manipulations to evaluate function. For example, efficiency of CRISPRi [75] targeting drops off sharply as a function of distance from the TSS [76]. Moreover, effective interpretation of experiments involving insertion of a strong polyadenylation sequence to make an RNA-level knockout [33, 77] relies on accurate knowledge of the TSS to minimize residual RNA fragment size. In our study, direct comparison of short- and long-read alignments over known genes and curated transcription start and termination sites exposed informative differences between the two platforms. The ends of the transcripts are much better represented in the long-read alignments than in the short-read alignments. We reason that this difference may arise due to multiple factors such as incomplete cDNA synthesis from random hexamer primers on fragmented RNA and heterogenous mapping of segmented short reads. In contrast, full-length cDNA synthesis on unfragmented RNA using a 3’-based oligoT primer and potential sequencing of the entire cDNA culminate in end-to-end coverage of transcripts in the long-read data set. This provides a qualitative advantage to the long-read RNA-seq for full-length characterization of transcripts.

### TASSEL improves integration of short- and long-read transcript assembly

Informed by the strengths and caveats of each of the two platforms, we developed a hybrid pipeline, TASSEL, to merge the short-read transcriptome with the stranded long-read transcriptome. We tested TASSEL against commonly used long-read transcript assembly methods such as FLAIR [53], Bambu [65] and StringTie Mix [29]. TASSEL had the lowest false negative rate of transcript assembly and unlike other methods, TASSEL led to assembly of complete transcripts on the correct strand. FLAIR was limited by assembly on the incorrect strand and incomplete assembly. Although StringTie Mix was able to assemble transcripts on the correct strand, it had a substantially higher false negative rate than TASSEL and assembled many transcripts with truncated ends. We speculate that StringTie Mix overweighs higher depth of short-read datasets that causes loss of long-read information. TASSEL also led to a substantial reduction in the transcripts that are assembled within an intron of the reference gene. The prime reason for the higher efficacy of TASSEL is the use of stranded long reads which not only resolves the conflicting orientation of neighboring reads but also enhances the coverage of correctly assembled transcripts. Additionally, as TASSEL is employed after alignment and assembly of short- and long-read transcripts, it is not biased towards short-read transcript depth and maintains long-read information. As a test of its advantage at the functional level, TASSEL had higher enrichment of known TSS, TTS, histone modification of active TSS, and RNA Pol II than StringTie Mix. Together, TASSEL outperforms contemporary assembly methods for assembly on the correct strand, complete end-to-end assembly, highest sensitivity of assembly, and unbiased integration of high depth short-read sequencing with the qualitative enrichment of long-read sequencing, which collectively culminates to superior manifestation of biologically relevant transcriptomic features.

We tested TASSEL on cheRNA transcripts which have been challenging to characterize due to their low abundance in cells, high variation from one cell type to another, and lack of canonical attributes of coding transcripts [37, 38]. In comparison to short- or long-read data alone or combined analysis with StringTie Mix, the use of TASSEL led to a marked improvement in the assembly of cheRNA. It corrected the segmentation of many cheRNA genes, resulting in enhanced definition of cheRNA transcription start sites as defined by RNA Pol II enrichment [68, 78]. As the majority of cheRNA are lncRNA, our TASSEL-based assembly marks the first successful use of a hybrid assembly method in enhanced detection as well assembly of lncRNA. It can therefore be used to resolve open questions about lncRNA such as their true molecular identity, convergence with other genomic as well transcriptomic features, and the role of antisense lncRNA. Collectively, our analyses of short- and long-read RNA-seq attributes and the development of stranding and merging pipelines can inform and improve future long-read transcriptome analyses beyond the cheRNA transcripts tested here.

## Materials and Methods Cell Culture

HAP1 cells were grown in IMDM media (Gibco, 12440-053), supplemented with 10% FBE (Seradigm, 3100-500) and 1% Pen/Strep (ThermoFisher 15140-122) to 80-90% confluency in T75 flasks at 37°C. HL1 mouse cardiomyocytes (#SCC065) were grown per manufacturer instructions in supplemented Claycomb Medium (51800C-500ML) + 10% FBS (TMS-016) + 0.1mM Norepinephrine with 30mM L-ascorbic acid (A0937, A7506) + 5% L-Glutamine (G7513) in 0.1% gelatin-coated (SF008) T75 flasks.

To prepare the cells for RNA harvesting, they were washed once with 1X PBS, detached with TrypLE Express (Gibco #12605010), quenched with FBE-containing media, then washed with PBS, and pelleted by centrifugation for 5 min at 250 x g. Cell pellets were either snap frozen with liquid nitrogen or processed immediately.

### Nuclear fractionation and RNA isolation

Cell pellets were resuspended in Buffer A (10 mM HEPES•KOH pH 7.5, 10 mM KCl, 10% glycerol, 340 mM sucrose, 4mM MgCl_2_, 1 mM DTT, 1 x Protease Inhibitor Cocktail (PIC) [1mM AEBSF, 0.8 μM aprotinin, 20 μM leupeptin, 15 μM pepstatin A, 40 μM bestatin, 15 μM E-64; from 200x DMSO stock]). Nuclear fractions were separated as described previously [37, 38]. 1 μL of a 1:10 diluted ERCC RNA Spike-in standards aliquot (Life Technologies 4456740) was added to each nuclear fraction prior to addition of TRIzol reagent (Life Technologies 15596026). RNA from each fraction was extracted as described previously [37, 38]. RNA was processed using Zymo RNA Clean & Concentrator kit (Zymo Research R1017) with on-column DNase digestion. Ribosomal RNA was removed using the Ribo-zero Gold rRNA depletion kit (Illumina MRZG12324) for HL1 short-read RNA-seq samples and the RiboMinus kit (Invitrogen A15026) for HL1 long-read RNA-seq as well as HAP1 short- and long-read samples.

### Library preparation for short-read RNA-seq

100 ng of rRNA-depleted RNA from each nuclear fraction was used with the NEBNext Ultra II Directional library kit (NEB E7765S) to prepare RNA-seq libraries. RNA-seq libraries of nuclear fractions (chromatin pellet extract and soluble nuclear extract) from three independent cultures were sequenced through the University of Chicago Genomics Core Facility on the Illumina HiSeq 4000 to obtain 50 bp single end reads.

### Library prep for long-read RNA-seq

Two HAP1 and two HL1 long-read sequencing libraries were generated as follows. RNA from each fraction was extracted with Zymo RNA Clean & Concentrator (R1019) and subjected to qPCR analysis (primer sequences in Supplementary Table 2) for confirmation of enrichment for respective fractions (see Figure S1A). Next, the chromatin fraction RNA was depleted for ribosomal RNA with the RiboMinus Eukaryote Kit v2 (A15020) and precipitated with ethanol, then polyadenylated with E. coli Poly(A) Polymerase (M0276S) (for later priming with long read sequencing primers) and precipitated with ethanol. Successful rRNA depletion was assessed using BioAnalyzer. Finally, rRNA-depleted and polyadenylated chromatin RNA was subjected to library preparation according to the Oxford Nanopore direct cDNA sequencing kit protocol (SQK:DCS109). Libraries were checked for purity and size on a TapeStation with genomic DNA reagents (5067-5366) and tape (5067-5365). 10-50 fmol of library was loaded onto a MinION (MIN-101B) flow cell (R9.4.1; FLO-MIN106D) and run for 48 hours. Real-time basecalling information and FASTQ files were generated with the MinKNOW software using default settings (HAP1 rep1: minQ 7; HAP1 rep2: minQ 7; HL1 rep1: minQ 7; HL1 rep2: minQ 9).

### Alignment of short reads

Read quality and adapter content was checked using FastQC [79]. All short-read RNA seq libraries passed the quality check. Reads were then aligned to reference genomes (hg38 for HAP1 and mm10 for HL1 reads, catenated with ERCC sequences) using hisat2 version 2.1.0 [80] with --rna-strandness R --dta options. Sam files were converted to sorted bam files using samtools [81].

### Long read processing and ERCC analysis

FASTQ files generated by the MinKNOW software for each cell line were concatenated and mapped either to the hg38+ERCC (HAP1) or mm10+ERCC (HL1) genomes using minimap2 [46] without canonical splice sites (-ax splice -un). For stranding analyses, FASTQ files were subjected to our stranding pipeline prior to mapping (see Stranding methodology section below). For coverage analyses between replicates, HL1 replicate 1 was further filtered post-run to have a minimum phred score of 9 to match the minimum phred score of 9 used for replicate 2 using NanoFilt (-q 9). After sorting and indexing the BAM files (samtools), ERCC counts were computed using BEDTools coverage (Quinlan, 2014) and averaged between replicates; transcripts with count averages above 0 were kept. Then, coverage analyses were performed using Stringtie2 [56] with options -A to generate gene coverage in addition to transcripts and -L to specify long reads. Replicates were then analyzed in tandem using the Stringtie --merge function with expression estimation mode on (-e) to create a reference GTF with the union of transcripts between both replicates and subsequently re-estimate coverage for both genes and transcripts in individual replicates. Tags per million (TPM) was used for analysis of coverage. For the ERCC graphs, coverage was plotted against amount of ERCC added, length, and fraction of nucleotides mapped. Coverage of ERCCs was analyzed between long and short read datasets by first averaging ERCC coverage between replicates (excluding any transcripts with coverage of 0), and then plotting the averaged replicate coverages between long and short read datasets. Coverage analysis for all transcripts between long and short read datasets was performed by re-estimating coverage between the replicates with Stringtie2, then re-estimating coverage between long and short read re-estimated datasets and plotting the TPMs. As a supplementary comparison analysis, BEDTools counts vs StringTie TPMs were also plotted against each other for ERCCs.

### Correlation of alignment of reads

For testing the correlation of alignment, bam files were converted to counts per million (CPM)-normalized bigwig files using the bamCoverage function of deepTools (-bs 1 --normalizeUsing CPM) [82]. Then, the multiBigwigSummary function of deepTools was used to compute correlation of coverages of aligned reads at all genomic regions binned at 1 kb region (-bs 1000). Correlation between the replicates was then plotted using plotCorrelation function (-c pearson -p scatterplot --log1p).

### Metagene plots

To plot the aligned short and long reads on the known genes: Coordinates of the known genes (mm10vm23 assembly for mouse genome and hg38v41 assembly for human genome) were obtained from the UCSC table browser [83]. Counts per Million (CPM)-normalized bigwig files (binned at 1 base pair) of aligned reads were then used to score mean coverage over the known genes using the computeMatrix function of deepTools in scale-regions mode (-b 1000 -a 1000 --missingDataAsZero) and plotted using plotHeatmap.

To plot the aligned short and long reads on transcription start sites and polyA sites: Coordinates of annotated transcription start sites were obtained from refTSS [49] and polyA sites from polyASite [50]. Mean coverage over these coordinates were then calculated using the reference-point mode of computeMatrix (-b 1000 -a 1000 --missingDataAsZero) and plotted using plotHeatmap.

### Comparison of transcript assembly programs

For long-read transcript assembly, we compared FLAIR, StringTie *denovo* and StringTie guided approaches. To assemble transcripts with FLAIR, sorted bam files were converted to bed12 format using the bam2Bed12.py script of FLAIR. FLAIR correct was used with reference genome fasta and annotation files (flair correct –nvrna -q input.bed -g refgenome.fa -f refgenome.gtf -o FLAIRcorrect_output.bed). The corrected bed file was then used to obtain a gtf file using FLAIR collapse (flair collapse -g refgenome.fa -q FLAIR_corrected.bed -f refgenome.gtf -r sample.fq -o flair_collapse_output.gtf). For StringTie, sorted bam files for each sample were used to assemble transcripts using StringTie version 2.1.1 [52] with the -L option in either *de novo* or guided assembly mode (-G reference_annotation.gtf). For guided assembly, reference annotation gtf files of the gencode hg38v35 assembly for HAP1 samples and the mm10 assembly for HL1 samples were concatenated with gtf files for the ERCC spike-in RNA mix.

For short-read transcript assembly, sorted bam files for each sample were used to assemble transcripts using cufflinks version 2.2.1 (-u -N --library-type fr-firststrand) [84] or StringTie version 2.1.1 (--rf) [52] in either *de novo* or guided assembly mode. In cufflinks guided assembly mode, reference genome transcripts that are not detected in the sample are appended to the sample gft files with their coverage marked as “0.0000”. As it would artifactually inflate the sensitivity of assembly, we removed all transcripts with zero coverage in the sample gtf files.

To compare the efficacy of transcript assembly programs, gtf files of transcripts assembled by each of the programs was compared with corresponding reference genome gtf files using gffcompare [85]. Sensitivity (*True positive / (True positive + False negative)*) and Precision (*True positive / (True positive + False positive)*) of transcript assembly were compared to gauge the efficacy of assemblers.

### Testing stranding with UNAGI

UNAGI [86] was used in the default mode (-i long-read.fastq -g ref_genome.fa). It yields a fastq file with stranded reads. Number reads in this stranded fastq file was used to calculate percent stranding.

### SLURP (Stranding Long Unstranded Reads using Primers) stranding methodology

We used MEME analysis [62] to predict the sequences of primers used in the cDNA library prep. These primer sequences were then used as guides to ascertain the strand of origin of long reads. For this, the following stranding methodology was adapted:

1. Reads with partial primer 1 sequence in the first 100 bp of reads, permitting 2 mismatches, were extracted using:

seqkit grep -s -i -P -R 1:100 -m 2 -p GCTCTATCTTCTTT

2. Reads with the reverse complement of partial primer 2 in the last 100 bp, permitting 2 mismatches, were extracted using:

seqkit grep -s -i -P -R -100:-1 -m 2 -p CCCAGCAATATCAG.

3. Reads from 1 and 2 were combined and deduplicated.

4. Reads with partial primer 2 sequence in the first 100 bp of reads, permitting 2 mismatches, were extracted using:

seqkit grep -s -i -P -R 1:100 -m 2 -p CTGATATTGCTGGG.

5. Reverse complement of reads from (4) were made using:

seqkit seq -t dna -r -p.

6. Reads from (3) and (5) were combined and deduplicated to obtain stranded reads.

A bash script of this stranding pipeline is made available on at the GitHub repository (https://github.com/ka-inth-amoldeep/SLURP).

### Detection and analysis of the palindrome artifact in long-read sequencing

We extracted common reads which had primer 1 in their first 100 bp (seqkit grep -s -i -P -R 1:100 -p GCTCTATCTTCTTT) and the reverse complement of primer 1 in their last 100 bp (seqkit grep -s -i -P -R -100:-1 -p AAAGAAGATAGAGC).

These reads were then analyzed using RNAfold [87], as we reasoned that the secondary structure prediction of RNAfold could be used to detect potential palindromes in long reads. From the RNA-fold dot and bracket output (dot denotes unpaired residue and bracket denotes paired residue), we calculated the percentage of brackets in the given read to determine extent of residues base-paired in the long-reads. In addition, RNA-fold also provides minimum free energy (MFE) scores of residues in the reads. We binned and scaled MFE scores of multiple reads to 100 bp to obtain a meta-mountain plot of MFE scores.

For mapping and counting statistics, the palindromic reads were aligned to the reference genome using minimap2 (-ax splice -un --MD). As palindromes were tagged with supplementary alignment to the same locus, we filtered out reads with the supplementary alignment sam tag using samtools (view -F 2048). The filtered and non-filtered bam files were then used to assemble and quantify transcripts using StringTie to compare coverage and abundance (TPM) of transcripts.

### Comparison of short- and long-read transcriptomes

For comparing abundance of transcripts and genes in short- and long-read samples, StringTie-assembled gtf files of long-read samples (2 replicates) and short-read samples (3 replicates) were merged using stringtie – merge. This merged transcriptome gtf file was then used as a reference for re-estimation of transcripts in each of the samples using stringtie -e -B -G stringtie_merge_file.gtf -C coverage_file -A gene_abundance_file. Then, average transcript and gene abundance (Tags per million, TPM) of every transcript and gene was calculated across the replicates for short-read and long-read samples. These average TPMs were compared to test the correlation between short-read sequencing and long-read sequencing.

To compare the transcript structure assembled by short-read and long-read assembly, gffcompare was used with the gtf file of long-read assembly as a reference transcriptome and the gtf file of short-read assembly as the query transcriptome. The relationship between the reference and query transcripts was ascertained by the “class code” output of gffcompare.

### Enrichment of transcript ends in short and long reads

CPM-normalized bedgraph files were made from sorted bam files of each sample using the bamCoverage function of deepTools (-bs 1 --normalizeUsing CPM -of bedgraph). Then, sorted bedgraph files were used to calculate mean signal at the 100 bp region around the TSS or TTS of known genes using bedtools [55] (map -a TSS_100bp.bed -b CPMnormlized.bg -c 4 -o mean > bedtoolsmapped_TSS100bp_mean). The average signal thus obtained was then normalized to the average signal obtained from mapping reads to a similar number of random regions (100 bp wide) in the genome.

### Long-read gene fusion

Name-sorted bam files of each long-read sample were used as input to detect fusion events using LongGF [88] with <min-overlap-len> 100 <bin_size> 30 <min-map-len> 100. Then, grep “SumGF” LongGF.log > FusionList was used to get a list of detected fusions. The list of fusion events was then used to make a circos plot using BioCircos [89].

### TASSEL (Transcript Assembly using Short and Strand-Emended Long reads)

Long reads were stranded using SLURP (described above). Stranded long reads and short reads were aligned to the reference genome using minimap2 and hisat2, respectively. StringTie2 was used to assemble transcripts in guided assembly mode (with -L option for long reads). Assembled transcripts were merged in a strand-aware manner using stringtie --merge --rf.

### Comparison of TASSEL with other assembly programs

TASSEL was compared to FLAIR, Bambu and StringTie Mix. For assembly with FLAIR, sorted bam files of combined replicates were converted to bed12 format using the bam2Bed12.py script of FLAIR. FLAIR correct was used with reference genome fasta and annotation files (flair correct --nvrna -q input.bed -g refgenome.fa -f refgenome.gtf -o FLAIRcorrect_output.bed). The corrected bed file was then used to obtain a gtf file using FLAIR collapse (flair collapse -g refgenome.fa -q FLAIR_corrected.bed -f refgenome.gtf -r sample.fq -o flair_ collapse_output.gtf).

Bambu (version 3.1.1) was obtained from github/goekelab. Annotation file for Bambu was prepared using BambuAnnotations < prepareAnnotations(ERCC_hg38gencodev35.gtf). Long-read alignments of combined replicates were obtained from minimap2 and were used to make a summarized experiment object using longread_Bambu_se <-Bambu(reads = longread.bam, annotations = BambuAnnotations, genome = ERCC_ hg38.fa). Transcripts assembled by Bambu were extracted using Bambu_constructedAnnotations = longread_ Bambu_se [assays(longread_Bambu_se)$fullLengthCounts > 0].

For StringTie Mix, bam files of combined short-read replicates and long reads replicates were obtained using hisat2 and minimap2, respectively. Reference annotation gtf files of the gencode hg38v35 assembly for HAP1 samples and the mm10 assembly for HL1 samples were concatenated with gtf files for the ERCC spike-in RNA mix. These files were then used with StringTie Mix (stringtie --mix short-read.bam long-read.bam –o stringtiemix.gtf -C stringtiemix -G ERCC_reference.gtf).

### Enrichment of H3K4me3, RNA Pol II and PRO-seq at TSS

Fastq files for H3K4me3 ChIP-seq in HAP1 cells (Input: SRR6671609, H3K4me3: SRR6671589) and RNA Pol II ChIP-seq for HAP1 cells (SRR2301045) [90] were downloaded from the GEO database. Reads were aligned to genome (hg38) using bowtie2 [91]. For H3K4me3, an input normalized bigwig file was made using the bamCompare function of deepTools (bamCompare -bs 1 -b1 H3K4me3_IP.bam -b2 Input.bam –operation ratio). For Pol II ChIP-seq, sorted bam files were directly converted to counts-normalized bigwig files using the bamCoverage function of deepTools (-normalizeUsing CPM). The bigwig files were then used to plot metagene profiles on TSSs of transcripts from StringTie Mix and TASSEL genes using the computeMatrix and plotProfile functions of deepTools.

For PRO-seq, bigwig files of normalized PRO-seq signal in HAP1 cells were obtained from GSM5829596 [92]. Forward PRO-seq signals were contoured over plus strand TSS and reverse PRO-seq signals were contoured over negative strand TSS of TASSEL or StringTie Mix transcripts using the computeMatrix function and plotted using the plotProfile function of deepTools.

### Estimation of cheRNA

StringTie and DESeq2 [93] were used to detect transcripts and genes which were significantly enriched in the chromatin fraction as compared to the nucleoplasm fraction in short-read sequencing. For this, gtf files of chromatin fraction reads and nucleoplasm fraction reads (three biological replicates each) were obtained by StringTie guided assembly of aligned reads. These gtf files were merged using stringtie --merge --rf. This merged transcriptome gtf file was then used as a reference for re-estimation of transcripts in each of the samples using stringtie -e -B -G stringtie_merge.gtf. Raw counts of transcript and gene abundance were obtained using the prepDE.py script from StringTie. The raw counts were then used to calculate differentially expressed transcripts and genes in the chromatin vs nucleoplasm fractions using DESeq2. Transcripts and genes which were enriched more than four-fold in the chromatin fraction at adjusted *p* < 0.05 and had a length of more than 200 bp were deemed as cheRNA.

### Gene ontology for cheRNA proximal genes

Protein coding genes closest to cheRNA genes were computed using the closestBed function of bedtools (-a cheRNA_genes -b refgenome_proteincodinggene). The identified genes were then uploaded to the DAVID gene ontology portal [94] to calculate enriched biological processes categories.

### Comparison and merge of cheRNA with long-read sequencing

To plot long-read alignments on cheRNA genes: CPM-normalized bigwig files of long-read alignments were used to calculate mean coverage over cheRNA genes using the computeMatrix function of deepTools in scale-regions mode (-b 1000 -a 1000 --missingDataAsZero) and plotted using plotHeatmap.

To calculate the extent of segmentation of cheRNA genes, genes in TASSEL-merged gtf file were further consolidated using bedtools merge (-s -c 4,6 -o distinct). Then we intersected these consolidated genes with cheRNA genes using intersectBed (-wa -wb -s) to obtain the number of consolidated genes that intersected more than once with different cheRNA genes, essentially counting the segmented cheRNA genes.

### Pol II occupancy at cheRNA

Fastq files for RNA Pol II ChIP-seq for HAP1 cells (SRR2301045) [90] and eight-week old mouse heart (RNA Pol II IP: SRR489670; Input: SRR489681) [95] were downloaded from the GEO database. Reads were aligned to respective genomes (hg38/mm10) using bowtie2 [91]. For HAP1 Pol II ChIP-seq, sorted bam files were directly converted to bigwig files using the bamCoverage function of deepTools. For mouse Pol II, an input normalized bigwig file was made using the bamCompare function of deepTools (bamCompare -bs 1 -b1 Pol_IP-.bam -b2 Input.bam –operation ratio). The bigwig files were then used to plot metagene profiles on TSSs of cheRNA genes using the plotProfile function.

## Availability of data and materials

Short- and long-read data generated for this study have been deposited at the Gene Expression Omnibus (GEO) under accession number GSE215355 and GSE215357. A standard script for SLURP and TASSEL is provided at the GitHub repositories (https://github.com/kainth-amoldeep/SLURP and https://github.com/ka-inth-amoldeep/TASSEL).

## Competing interests

The authors declare that they have no competing interests.

## Funding

G.A.H. is supported by the NIH Genetics & Regulation Training Grant (T32 GM07197) and the Genetic Mechanisms and Evolution Training Grant (T32 GM139782). This work is supported by NIH grants (R01HL148719 and R35GM145373) to A.J.R.

## Authors’ contributions

G.A.H. and A.J.R. conceived of the study. A.S.K. made short-read RNA-seq libraries for HAP1 cells. J.M.H. made short-read RNA-seq libraries for HL1 cells. G.A.H. made long-read RNA-seq libraries for HAP1 and HL1 cells. A.S.K and G.A.H. performed computational analyses with input and oversight from A.J.R. A.S.K. and G.A.H. wrote the manuscript with input from A.J.R. All authors read and approved the final manuscript.

## Acknowledgements

We thank Peter Faber, Lindsay Scarpitta, and Mikayla Marchuk in the University of Chicago Functional Genomics Facility for Illumina sequencing. We thank members of the Ruthenburg and Ivan Moskowitz labs at the University of Chicago for helpful suggestions to improve the manuscript.

## Notes

### Competing Interest Statement

The authors have declared no competing interest.

