## Supplementary Information for "Merging short and stranded long reads improves transcript assembly"

Supplementary note 1

Supplementary figures (S1-S6)

Supplementary tables (1-2)

Supplementary references

**Supplementary note 1: Long-read alignments capture novel cell line-specific gene fusion events**

An important feature that can be detected with long reads is gene fusion. Long reads are particularly useful in the reliable detection of gene fusions, as they not only detect the junction but also are more likely to cover uniquely mappable sequences on both sides of the discontinuity. Using LongGF [1] with long reads from HAP1 cells, we detected >20 gene fusion events supported by at least two reads in each of the two HAP1 long-read replicates (S1I Fig). Notably, we were able to detect the BCR::ABL fusion which is a key mutagenic fusion in this leukemia cell line [2]. We compared gene fusion events in our long-read transcriptome data with a comparable long-read genomic DNA dataset in HAP1 cells [3]. Surprisingly, we found only one common fusion, which was BCR::ABL. Therefore, our long-read transcriptome analyses unveiled novel gene fusion events in the HAP1 cells, possibly due to enhancement of sensitivity in our chromatin-focused pool of transcripts. Intriguingly, we did not detect any gene-fusion in the non-leukemia HL1 cells (S1I Fig), suggesting a cell line-specific detection of translocations in our dataset that is not due to technical artifacts in the library preparation.

**Figure S1. Supporting data for Figure 1**

**A** RT-qPCR confirmation of RNA fractionation into chromatin (purple) and nucleoplasm (green) extracts in the indicated samples. Protein coding transcripts (*GAPDH* and *B2M*) were used as representative of nucleoplasm enrichment while lncRNA (*MALAT1* and *PVT1*) were used as controls for chromatin enrichment. Error bar indicates SD for replicates (n=3 for short-read; n=2 for long-read).

**B** Density plot of read lengths from long-read sequencing of HL1 replicates (n = 2).

**C** Correlation between HL1 replicates (n = 2) for genomic coverage of long-read alignments binned at 1 kb. Pearson correlation is shown.

**D** Correlation between HAP1 (left) and HL1 (right) replicates (n = 3) for genomic coverage of short-read alignments (*top* nucleoplasm fraction; *bottom* chromatin fraction) binned at 1 kb. Pearson correlations are shown for each plot.

**E** Average counts per million (average CPM, dark color; +/- SD, lighter color) of aligned reads across all ERCC transcripts, meta-scaled to 1000 nt, in the HL1 short-read (n=3) and long-read (n=2) samples.

**F** Relationship between the average fraction of ERCC transcripts covered by short (left) or long (right) read alignments as a function of their amount (attomoles) or length (nt) for HL1 samples.

**G** Metagene plots and heatmaps of mapped read coverage (average CPM) for HL1 short- and long-read alignments, scaled to transcription start sites (TSS) and transcription termination sites (TTS) of known genes.

**H** Metagene plots and heatmaps of mapped read coverage (average CPM) for HL1 short (top) and long (bottom) reads, centered on curated transcription start sites (TSS; [4]) or termination sites (polyA; [5]).

**I** Circos plot depicting gene fusions supported by at least 2 reads in HAP1 (top) and HL1 (bottom) long-read datasets. Dark blue arcs indicate fusion events detected in both replicates; light blue arcs indicate fusion events detected in only one of the two replicates. The BCR::ABL fusion, a key genomic fusion, is highlighted in red.

### Figure S1

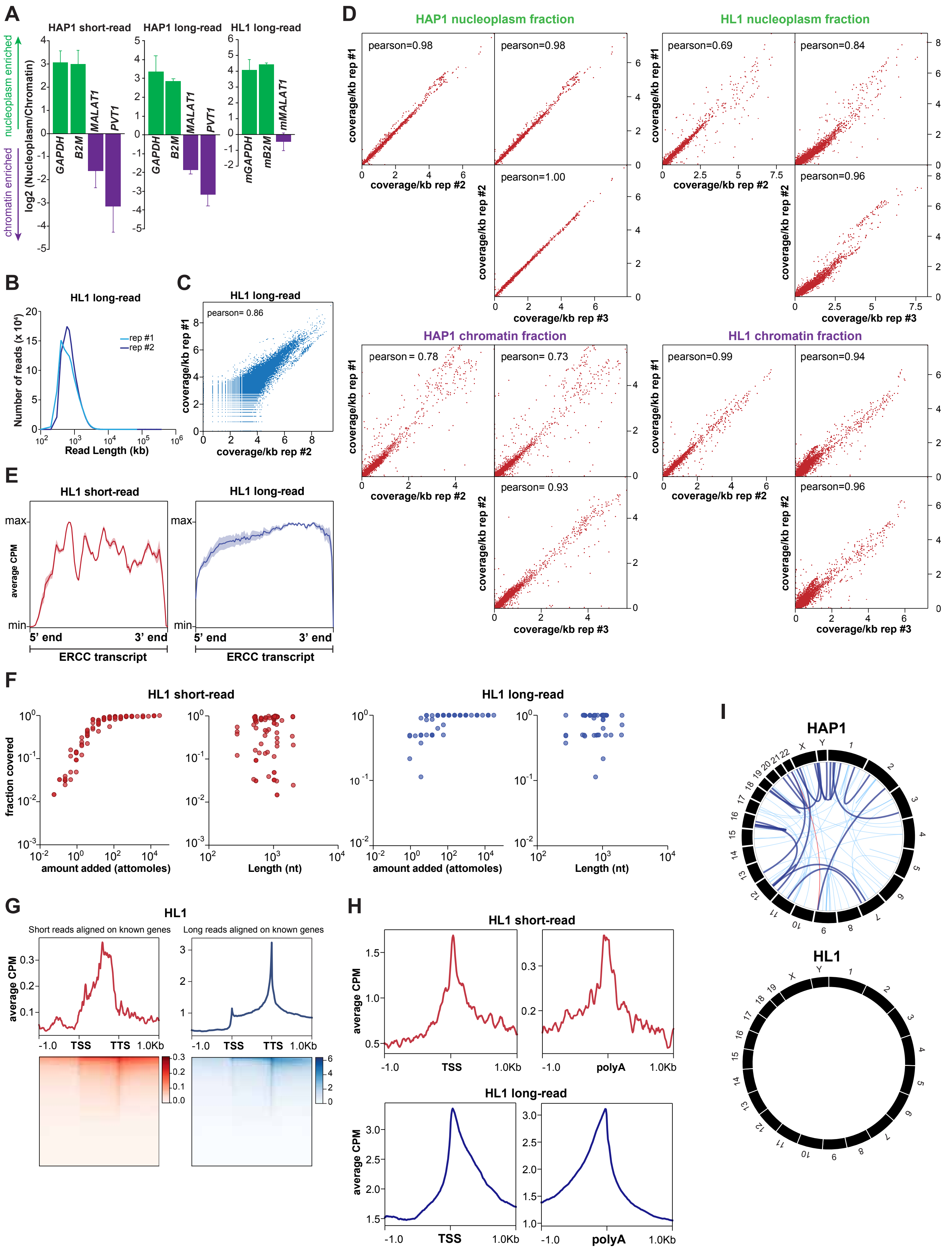

A
HAP1 short-read
HAP1 long-read
HL1 long-read
nucleoplasm enriched
chromatin enriched
log2 (Nucleoplasm/Chromatin)
GAPDH
B2M
MALAT1
PVT1
mGAPDH
mB2M
mMALAT1
B
HL1 long-read
Number of reads (x 10³)
rep #1
rep #2
Read Length (kb)
C
HL1 long-read
pearson= 0.86
coverage/kb rep #1
coverage/kb rep #2
D
HAP1 nucleoplasm fraction
HL1 nucleoplasm fraction
HAP1 chromatin fraction
HL1 chromatin fraction
pearson=0.98
pearson=0.98
pearson=1.00
pearson=0.69
pearson=0.84
pearson=0.96
pearson = 0.78
pearson= 0.73
pearson= 0.93
pearson=0.99
pearson=0.94
pearson=0.96
coverage/kb rep #1
coverage/kb rep #2
coverage/kb rep #3
E
HL1 short-read
HL1 long-read
average CPM
max
min
5' end
3' end
ERCC transcript
F
HL1 short-read
HL1 long-read
fraction covered
amount added (attomoles)
Length (nt)
G
HL1
Short reads aligned on known genes
Long reads aligned on known genes
average CPM
TSS
TTS
1.0Kb
H
HL1 short-read
HL1 long-read
average CPM
TSS
polyA
1.0Kb
I
HAP1
HL1

**Figure S2. Supporting data for Figure 2**

**A** Comparison of the indicated short-read (left) and long-read (right) transcript assembly methods. Shown are the sensitivity and precision of assembly at the transcript level as deduced by comparing assembled transcripts of HAP1 chromatin fraction replicates (n=3) with the reference genome transcripts using gffcompare. Error bar indicates SD.

**B** Average abundance (Tags Per Million; TPM) of ERCC transcripts determined by StringTie versus amount added (attomoles) or length (nt) of the transcript in HL1 samples. (n=3 for short reads, n=2 for long reads). Short-read plots are shown in red (left), long-read plots in blue (right). $R^2$, correlation coefficient for linear regression.

**C** Comparison of abundance (TPM) of ERCC transcripts obtained from StringTie with BEDTools counts for HL1 long-read replicate 1. Both methods performed nearly identically ($R^2$=0.99, correlation coefficient for linear regression).

**D** Correlation between HL1 replicates for transcriptome assembly by StringTie. Shown are the scatter plots of abundance (TPM) of each transcript in the two replicates. Colors correspond to kernel density estimations of scatter plot distribution. $R^2$, correlation coefficient for linear regression.

**E** Comparison of structure of transcripts assembled by StringTie for HAP1 and HL1 short-read samples with structure of reference genome transcripts. The class codes for relationship between the assembled transcript and the closest reference transcript were deduced from gffcompare.

**F** Correlation plots for the average abundance (TPM) of ERCC spike-in transcripts in the short-read datasets (n=3) with the average abundance in the long-read datasets (n=2) of HL1 samples, determined by StringTie. Shown are the transcripts detected by both sequencing platforms.

**G** Scatter plots comparing the average abundance (TPM) of all transcripts in the short-read datasets (n=3) and the long-read datasets (n=2) in HL1 samples. Abundance was calculated using the re-estimation function of StringTie from the merged transcriptome. Color bars correspond to kernel density estimations of scatter plot distribution.

**H** Histograms of length of transcripts assembled in the short- and long-read transcriptome. The distribution is shown for transcripts up to 10 kb in length.

### Figure S2

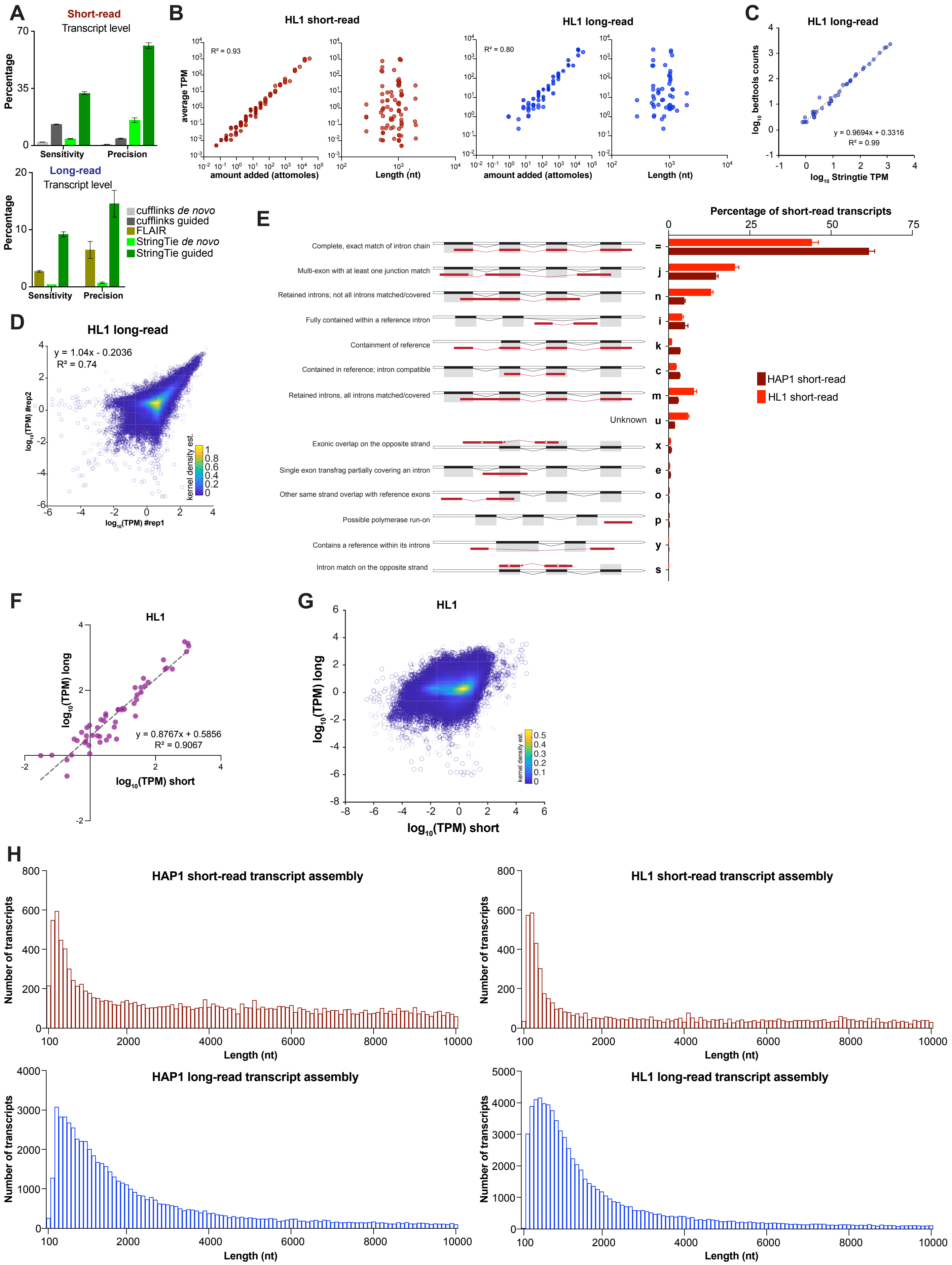

A
Short-read
Transcript level
Percentage
Sensitivity
Precision
Long-read
Transcript level
cufflinks de novo
cufflinks guided
FLAIR
StringTie de novo
StringTie guided
B
HL1 short-read
R² = 0.93
average TPM
amount added (attomoles)
Length (nt)
HL1 long-read
R² = 0.80
C
HL1 long-read
log10 bedtools counts
y = 0.9694x + 0.3316
R² = 0.99
log10 Stringtie TPM
D
HL1 long-read
y = 1.04x - 0.2036
R² = 0.74
log10(TPM) #rep2
log10(TPM) #rep1
kernel density est.
E
Percentage of short-read transcripts
Complete, exact match of intron chain
Multi-exon with at least one junction match
Retained introns; not all introns matched/covered
Fully contained within a reference intron
Containment of reference
Contained in reference; intron compatible
Retained introns, all introns matched/covered
Unknown
Exonic overlap on the opposite strand
Single exon transfrag partially covering an intron
Other same strand overlap with reference exons
Possible polymerase run-on
Contains a reference within its introns
Intron match on the opposite strand
= j n i k c m u x e o p y s
HAP1 short-read
HL1 short-read
F
HL1
log10(TPM) long
y = 0.8767x + 0.5856
R² = 0.9067
log10(TPM) short
G
HL1
log10(TPM) long
kernel density est.
log10(TPM) short
H
HAP1 short-read transcript assembly
HL1 short-read transcript assembly
HAP1 long-read transcript assembly
HL1 long-read transcript assembly
Number of transcripts
Length (nt)

**Figure S3. Supporting data for Figure 3**

**A** Percent of long reads stranded by UNAGI [6] and SLURP (Stranding Long Unstranded Reads using Primers) in HAP1 and HL1 replicates.

**B** Detection of primer 1 and primer 2 sequences in the long reads using MEME [7] *de novo* motif discovery.

**C** Measurement of indicated stranding methods in terms of the number of transcripts assembled and extent of transcripts mapping to the wrong strand of the reference genome. 3crit indicates using primer 1, primer 2 and reverse complement of primer 2; 4crit indicates primer 1, reverse complement of primer 1, primer 2 and reverse complement of primer 2; m2 indicates allowing 2 mismatches and m3 indicates allowing 3 mismatches.

**D** Workflow of SLURP pipeline. Reads that contain the first-strand synthesis primer (primer 1) or reverse complement (rc) of the strand switching primer (primer 2) are non-redundantly merged with the reverse complement of reads that contain the strand switching primer.

**E** Efficacy of stranding pipeline in a previously published dataset [8]. Shown are the percentage of transcripts mapping to the incorrect strand before and after stranding.

### Figure S3

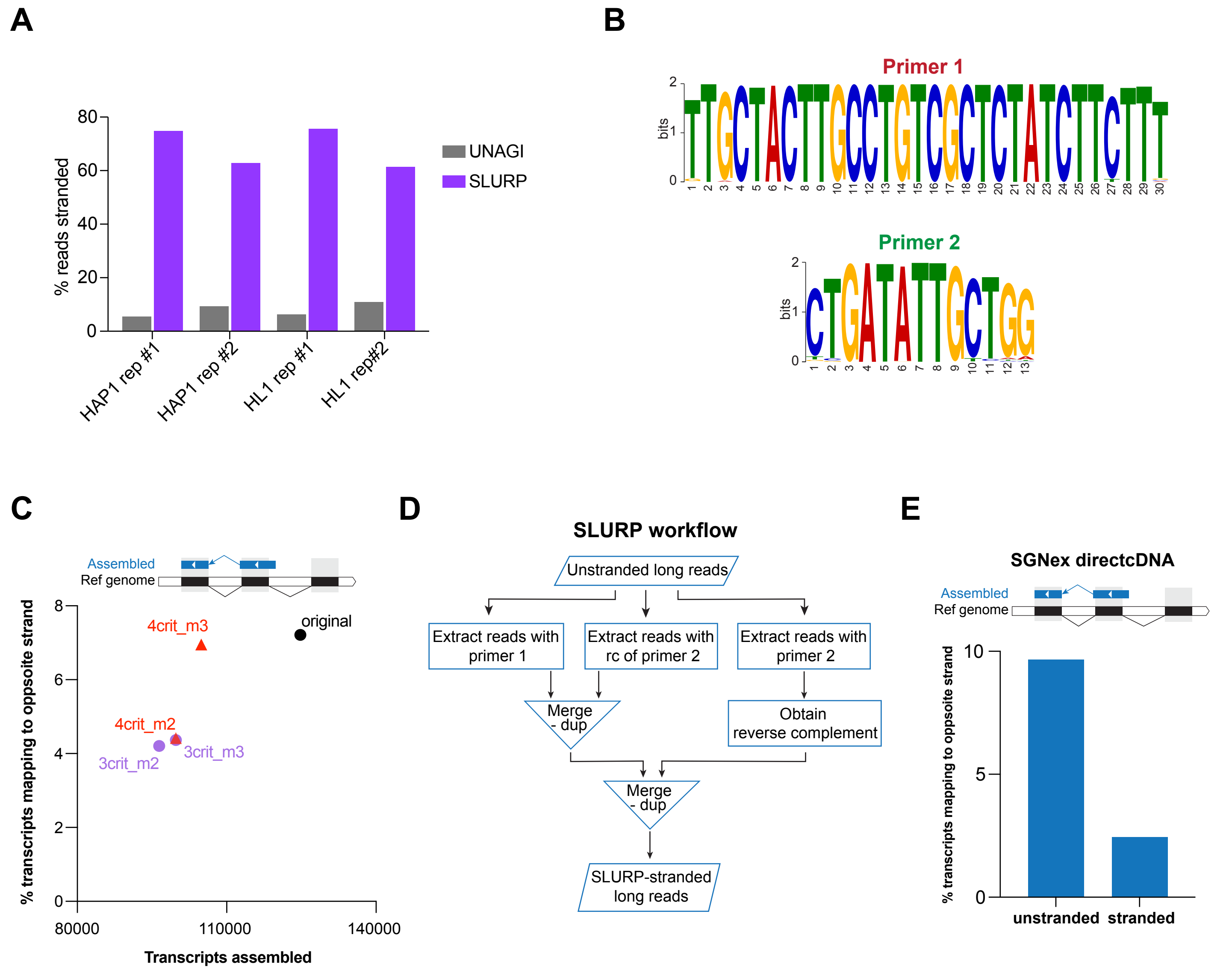
A
B
C
D
E
UNAGI
SLURP
% reads stranded
HAP1 rep #1
HAP1 rep #2
HL1 rep #1
HL1 rep#2
Primer 1
Primer 2
bits
Assembled
Ref genome
4crit_m3
original
4crit_m2
3crit_m3
3crit_m2
% transcripts mapping to oppsoite strand
Transcripts assembled
SLURP workflow
Unstranded long reads
Extract reads with primer 1
Extract reads with rc of primer 2
Extract reads with primer 2
Merge - dup
Obtain reverse complement
Merge - dup
SLURP-stranded long reads
SGNex directcDNA
unstranded
stranded

**Figure S4. Supporting data for Figure 4**

**A** Density plot of percentage of paired bases in the artifactual reads as a test for palindromes in the reads. RNAfold was used to determine paired bases in the secondary structure where a palindrome outweighs other structures.

### Figure S4

**A**

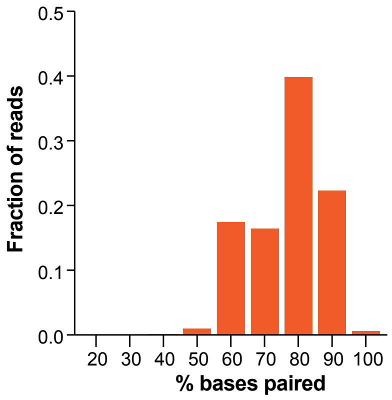

0.5
0.4
0.3
0.2
0.1
0.0
Fraction of reads
20 30 40 50 60 70 80 90 100
% bases paired

**Figure S5. Supporting data for Figure 5**

**A** Ends of the 92 ERCC transcripts (arranged in the increasing order of length) assembled by TASSEL (magenta circle) and StringTie Mix (StMix, green diamond) in HL1 dataset. Gray bar indicates actual transcript. The color bar indicates the abundance of the given transcript.

**B** Percent of assembled transcripts that match completely with a transcript (left) or contained within an intron (right) of reference transcript, using StringTie Mix or TASSEL in the HL1 dataset.

**C** Average H3K4me3 signal at the TSS assembled by StringTie Mix or TASSEL in the HAP1 dataset. Mean signal from TSS to +400 bp was computed for each of the TSS. Lower and upper whiskers mark $5^{th}$ and $95^{th}$ percentile, respectively. **** $p < 0.0001$, unpaired Student's t-test with Welch's correction.

#### Figure S5

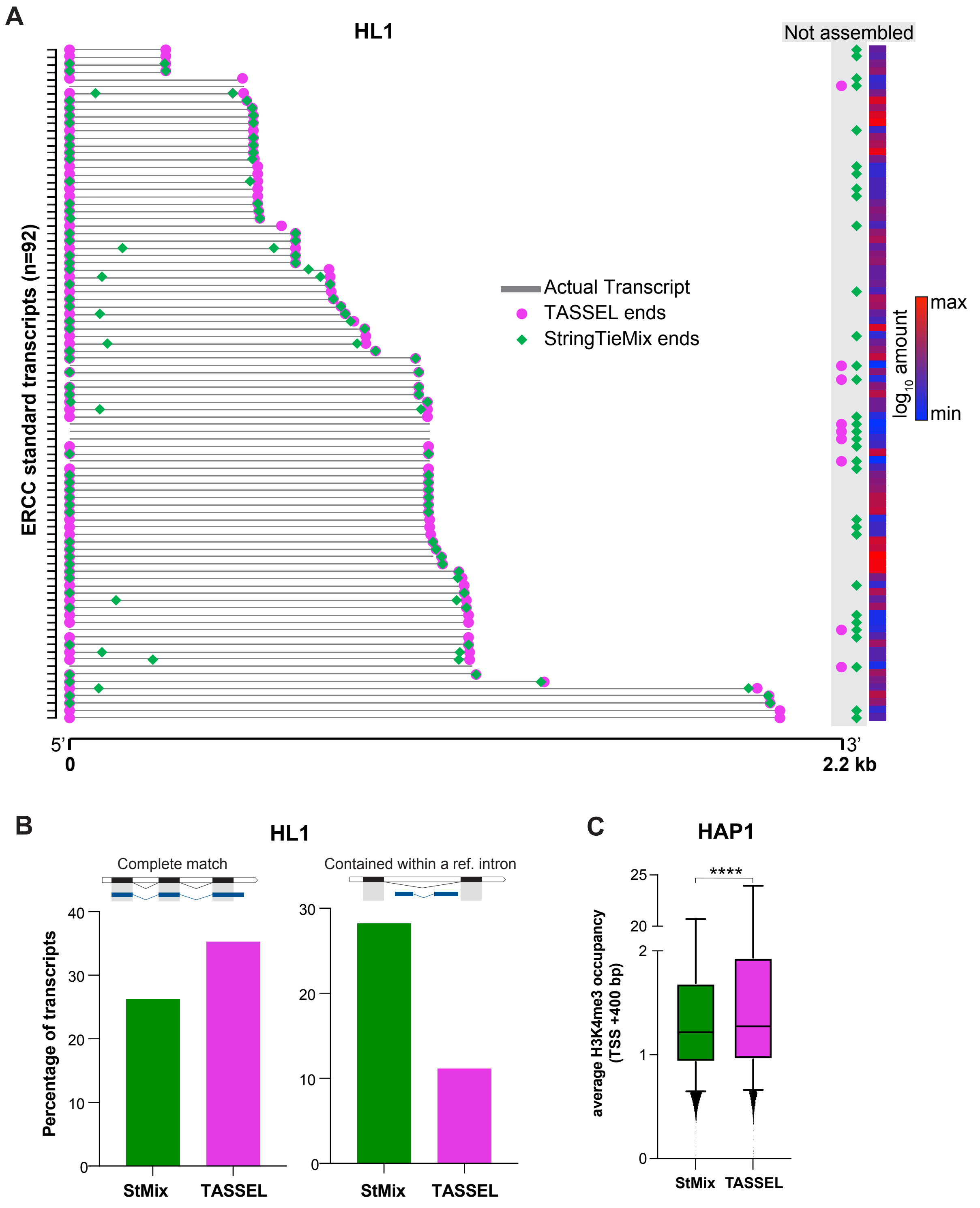
A
HL1
Not assembled
ERCC standard transcripts (n=92)
Actual Transcript
TASSEL ends
StringTieMix ends
max
min
log10 amount
5'
3'
0
2.2 kb
B
HL1
Complete match
Contained within a ref. intron
Percentage of transcripts
40
30
20
10
0
StMix
TASSEL
30
20
10
0
StMix
TASSEL
C
HAP1
****
25
20
2
1
0
average H3K4me3 occupancy (TSS +400 bp)
StMix
TASSEL

**Figure S6. Supporting data for Figure 6**

**A** Average count of genes in the chromatin and nucleoplasm fractions in HAP1 (left) and HL1 (right). Normalized gene counts obtained through DESeq2 were averaged for replicates (n=3).

**B** Gene ontology enrichment analyses of protein-coding genes closest to cheRNA genes detected in HAP1 and HL1 datasets. Top 15 categories detected by DAVID [9] and associated number of genes are shown. Color bars correspond to false discovery rate (FDR).

**C** Comparison of the abundance (normalized DESeq2 gene count) of cheRNA genes showing minimum (0-10%) and maximum (90-100%) overlap with a corresponding long-read gene. Data points outside 1.5x of Inter-quartile range were removed as outliers. **** $p < 0.0001$, Two-tailed Mann-Whitney test.

**D** Metagene plot depicting average RNA Pol II occupancy (solid lines; shaded regions indicate SE) at the TSS (±1Kb) of segmented, unsegmented and TASSEL-refined cheRNA genes in HL1 dataset. RNA Pol II occupancy calculated from ChIP-seq data from heart of an eight-week-old mouse [10].

### Figure S6

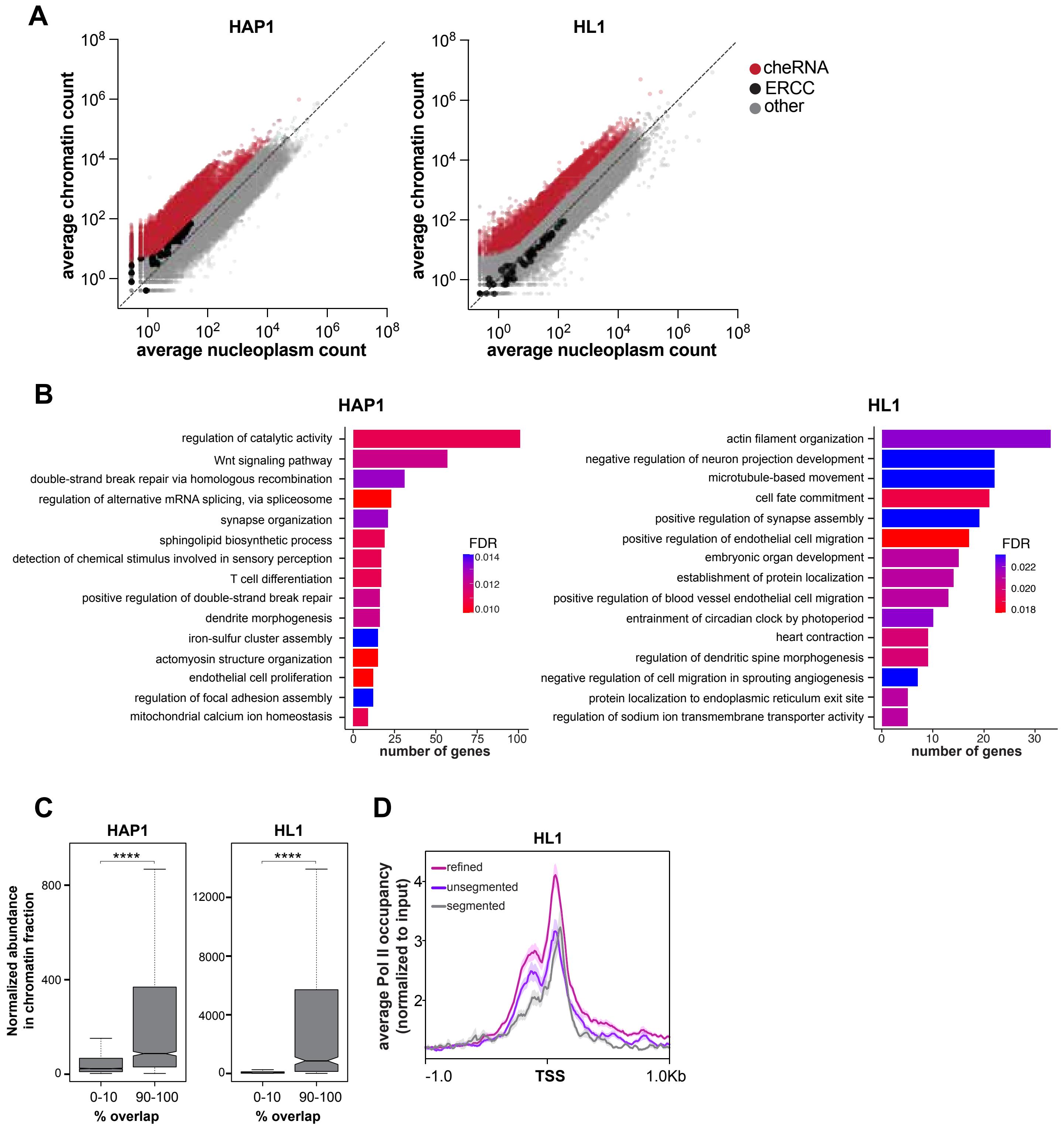
A
HAP1
HL1
average chromatin count
average nucleoplasm count
cheRNA
ERCC
other
B
HAP1
regulation of catalytic activity
Wnt signaling pathway
double-strand break repair via homologous recombination
regulation of alternative mRNA splicing, via spliceosome
synapse organization
sphingolipid biosynthetic process
detection of chemical stimulus involved in sensory perception
T cell differentiation
positive regulation of double-strand break repair
dendrite morphogenesis
iron-sulfur cluster assembly
actomyosin structure organization
endothelial cell proliferation
regulation of focal adhesion assembly
mitochondrial calcium ion homeostasis
FDR
number of genes
HL1
actin filament organization
negative regulation of neuron projection development
microtubule-based movement
cell fate commitment
positive regulation of synapse assembly
positive regulation of endothelial cell migration
embryonic organ development
establishment of protein localization
positive regulation of blood vessel endothelial cell migration
entrainment of circadian clock by photoperiod
heart contraction
regulation of dendritic spine morphogenesis
negative regulation of cell migration in sprouting angiogenesis
protein localization to endoplasmic reticulum exit site
regulation of sodium ion transmembrane transporter activity
FDR
number of genes
C
HAP1
HL1
Normalized abundance in chromatin fraction
****
0-10
90-100
% overlap
D
HL1
refined
unsegmented
segmented
average Pol II occupancy (normalized to input)
-1.0
TSS
1.0Kb

| Sample library | Reads generated | Reads passing QC | Reads mapped | Bases mapped | Median read length |
|---|---|---|---|---|---|
| HAP1 long-read chromatin fraction #1 | 1.24 million | 1.13 million | 96.84% | 90.83% | 1.97kb |
| HAP1 long-read chromatin fraction #2 | 2.94 million | 1.82 million | 99.03% | 90.06% | 1.7kb |
| HL1 long-read chromatin fraction #1 | 1.79 million | 1.55 million | 93.50% | 86.24% | 1.39kb |
| HL1 long-read chromatin fraction #2 | 2.5 million | 1.68 million | 96.51% | 85.26% | 1.26kb |
| HAP1 short-read chromatin fraction #1 | 127.27 million | 127.27 million | 96.67% | 96.43% | 51 nt |
| HAP1 short-read chromatin fraction #2 | 127.82 million | 127.82 million | 96.44% | 96.20% | 51 nt |
| HAP1 short-read chromatin fraction #3 | 129.68 million | 129.68 million | 96.54% | 96.29% | 51 nt |
| HAP1 short-read nucleoplasm fraction #1 | 126.03 million | 126.03 million | 97.64% | 97.36% | 51 nt |
| HAP1 short-read nucleoplasm fraction #2 | 126.85 million | 126.85 million | 97.70% | 97.42% | 51 nt |
| HAP1 short-read nucleoplasm fraction #3 | 131.55 million | 131.55 million | 97.73% | 97.46% | 51 nt |
| HL1 short-read chromatin fraction #1 | 354.05 million | 354.05 million | 87.10% | 86.81% | 51 nt |
| HL1 short-read chromatin fraction #2 | 337.90 million | 337.90 million | 83.88% | 83.56% | 51 nt |
| HL1 short-read chromatin fraction #3 | 327.32 million | 327.32 million | 85.37% | 85.05% | 51 nt |
| HL1 short-read nucleoplasm fraction #1 | 335.65 million | 335.65 million | 77.07% | 77.65% | 51 nt |
| HL1 short-read nucleoplasm fraction #2 | 327.32 million | 327.32 million | 65.43% | 65.02% | 51 nt |
| HL1 short-read nucleoplasm fraction #3 | 349.83 million | 349.83 million | 72.96% | 72.61% | 51 nt |

**Supplemental Table 1: Sequencing output of short- and long-read RNA-seq.**

| Primer name | Sequence |
| --- | --- |
| human_GAPDH F | CCGGGAAACTGTGGCGTGATGG |
| human_GAPDH R | AGGTGGAGGAGTGGGTGTCGCTGTT |
| human_B2M F | CTCTCTCTTTCTGGCCTGGAG |
| human_B2M R | TCTGCTGGATGACGTGAGTA |
| human_PVT1 F | CTGAGCCCCACTTCCTCTTG |
| human_PVT1 R | TCATCACGCTCCCCTAGCTT |
| human_MALAT1 F | GGAGCTTGAGGAAACCGCAGATAAG |
| human_MALAT1 R | GCTTCATCTCAACCTCCGTCATG |
| mouse_GAPDH F | CATCACTGCCACCCAGAAGACTG |
| mouse_GAPDH R | ATGCCAGTGAGCTTCCCGTTCAG |
| mouse_B2M F | CATGGCTCGCTCGGTGAC |
| mouse_B2M R | CAGTTCAGTATGTTCGGCTTCC |
| mouse_MALAT1 F | CGTTTGAAGGCATGAGTTGG |
| mouse_MALAT1 R | TGCCTCCCAAGTGCTAGGAT |

**Supplemental Table 2: Primers used for RT-qPCR analyses.**
